# Nonlinear dependency of the bacterial flagellar motor speed on proton motive force and its consequences for swimming

**DOI:** 10.1101/2024.10.07.617036

**Authors:** Ekaterina Krasnopeeva, Lucas Le Nagard, Uriel E. Barboza-Perez, Wilson Poon, Chien-Jung Lo, Teuta Pilizota

## Abstract

The bacterial flagellar motor enables bacteria to swim by rotating helical flagellar filaments that form a bundle at the back of the cell. *Escherichia coli*’s motor uses the energy stored in the electrochemical gradient of protons, the proton motive force (PMF), to generate the torque driving this rotation. Until now, motor speed was thought to be proportional to the PMF, irrespective of the viscous load on the motor, and across the physiological range of PMF values. Here, we show that the PMF-speed relationship is non-linear in the high-torque regime. Because saturation in the relationship occurs in the physiologically relevant range, it challenges all current models of motor function that assume a tight coupling between the motor rotation and PMF across the entire physiological range. Furthermore, we experimentally determine the load on the motor experienced by swimming cells, and show that free swimming occurs close to or within the saturation regime, making the observed limiting torque evolutionary relevant.

## Introduction

Many motile bacteria, such as *Escherichia coli* and *Salmonella enterica*, swim to search for favourable environments [1–5]. They use a bundle of rotating helical flagella, each driven by a bacterial flagellar motor (BFM). The BFM is a ∼ 45 nm in size nanomachine and one of only a few known examples of a biological rotary motor [6–10].

In the case of *E. coli*, the BFM consists of a rotor composed of the cytoplasmic and membranous/supramembranous rings, and of a stator made up of a varying number of MotA_5_MotB_2_ units [11–17]. Stator units produce the torque driving the motor rotation by allowing protons to cross the plasma membrane through MotB channels [6, 18]. This inward flux of protons is driven by the proton motive force (PMF), an electrochemical gradient of protons that metabol-ically active cells maintain across their inner membrane [19–22].

Within the stator, MotA_5_MotB_2_ complexes dynamically bind/unbind from the motor and remain bound for typically 30 s before returning to a pool of ∼ 100 units diffusing in the membrane [23]. The average number of torque-generating units bound to the BFM increases with the motor torque, which is influenced by the magnitude of the PMF and by the viscous load attached to the motor [13, 14, 24]. The motor exhibits a characteristic torque-speed curve with a high-torque/low-speed plateau followed by a low-torque/high-speed regime [6, 25–28].

Despite the above-mentioned mechanosensing characteristics, two seminal studies found that a remarkably simple relationship links the output of the motor to its driving force, the PMF [24, 29]. Those studies showed that the rotation speed of the BFM is proportional to the PMF across a range of PMF values, under both low and high viscous loads. A similar proportionality to sodium motive force (SMF) was observed for sodium-powered motors, although the chemical and electrical components of the SMF had non-equivalent contributions at low sodium concentrations and at low load for the chimeric SMF-driven *E. coli* motor [22, 30–33]. Consequently, all current models that describe motor function, including the most recent one, implement tight coupling between motor rotation and PMF throughout the operating range [34].

While the previous work spans a set of PMF values [24], it does not cover its full physiological range [35,36]. Given that there must be a physical limit to the speed and torque that the BFM can deliver, it is interesting to enquire whether this limit could be reached in physiologically relevant conditions, and what the consequences of such a limit would be for our understanding of the motor function and bacterial swimming. We therefore measured the rotational speed of the BFM across a wide range of physiologically achievable PMF values, increasing the PMF by providing a given amount of carbon source or decreasing it with the addition of ionophores or by inducing light damage, all while subjecting the motor to different viscous loads. By so exploring the entire PMF-torque space available to *E. coli* we find, contrary to previous thoughts, that the PMF-speed proportionality does not hold universally and saturates at higher PMF values and high viscous loads. We demonstrate that the relationship is limited by the maximum torque the motor can achieve. Moreover, we show that fully energised free-swimming cells can experience this limit in aqueous environments, where small increases in viscosity result in a swimming speed that is independent of the PMF. Combining these results, we experimentally determine the mechanical load under which flagellar motors of free-swimming *E. coli* operate. Finally, we discuss our results in the context of previous work and offer explanations for apparent discrepancies.

## Results

### The linear relationship between BFM speed and PMF fails at high load

To probe the limits of the PMF-speed proportionality, we first recorded the speed of individual motors, each on a separate cell, in the presence or absence of a carbon source. *E. coli* were grown in LB and washed in MM9 medium containing either no carbon source or 0.3% glucose, which is equivalent to 16.65 mM glucose and assumed to generate maximum PMF [37]. Thus, we obtained cells with moderate or high PMF, respectively. To record BFM speeds under different mechanical loads, we used the bead assay [3, 6, 26], whereby we attach polystyrene spheres (beads) of various diameters to the flagella stubs driven by the motor (see *Materials and Methods*).

Contrary to expectations, the relative speed increase induced by glucose supplementation varied depending on the bead size, Fig. 1A and Fig. SI 1. For the smaller loads (0.35 µm and 0.5 µm beads), glucose addition induced a ∼ 2-fold increase in BFM speed, while for higher loads (1 µm and 1.5 µm beads) the speed increased by only 10-20%. Those differences indicate that the PMF-speed proportionality is not independent of the load.

**Fig. 1.**
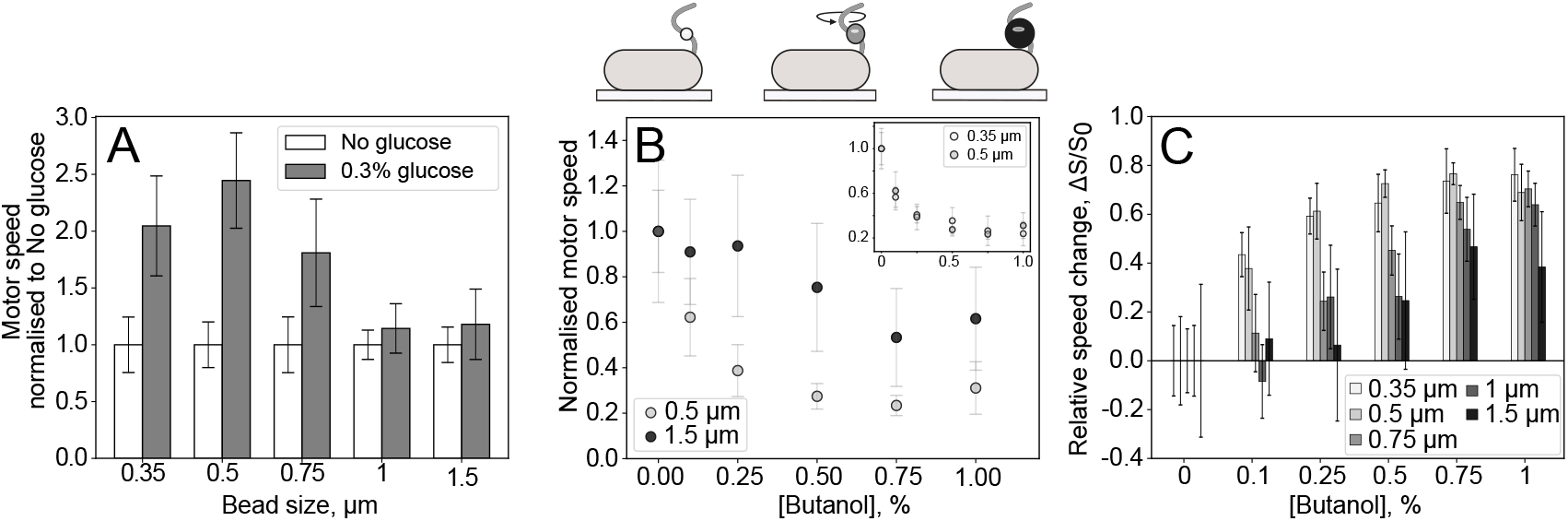
Relative changes in flagellar motor speed at different PMF values depend on the viscous load on the motor. The top gives a cartoon of the bead assay, with different beads scaled to size between each other (including colour scaling), but not with respect to the cell. **A**. Relative increase in BFM speed upon glucose addition depends on the viscous load. Rotational frequencies of the BFM are measured with and without glucose, and normalised to the speeds with no glucose, for beads of different sizes (0.35 µm to 1.5 µm). Glucose addition induces an approximately 2-fold speed increase for motors under low-load (0.35 µm, 0.5 µm beads), while those under high-load experience only a 20% speed increase. **B**. Load-dependent relative decrease in BFM speed upon butanol addition. BFM speed under 0.5 µm and 1.5 µm beads load is measured at different butanol concentrations and normalised by the 0% butanol speed. The BFM speed stays constant up to 0.25% butanol addition under high load, where the low load motor speed is already reduced by 60%. (***Inset***) 0.35 µm and 0.5 µm beads display identical speed decays upon butanol addition. **C**. BFM speed sensitivity to butanol scales inversely with the load. The change in BFM speed, Δ*S/S*_0_ = (*S*_0_ − *S*)*/S*_0_, where *S* is the speed at a given butanol concentration and *S*_0_ is the 0% butanol speed, is measured with beads of different sizes. Relative speed changes are less pronounced at higher loads for all concentrations. **A.-C**. Error bars are standard deviations and *Materials and Methods* contains normalisation procedures, and Table SI 1 sample sizes.

To test whether this observation was specific to glucose and to probe motor speed variations in a wider PMF range, we exposed cells to butanol. Looking at the speed changes of motors loaded with 0.5 µm beads, we previously demonstrated that butanol behaves as a (pore forming [38]) ionophore, collapsing *E. coli*’s PMF in a concentration-dependent manner [39]. Similarly, we exposed the cells washed in MM9+0.3% glucose to different concentrations of butanol (0-1%), while measuring the motor speed after butanol addition, but this time for different bead sizes.

Again, the relative changes in the BFM speed depended on the motor load, Fig. 1B, C, and Fig. SI 2. For 0.5 µm beads, the speed was inversely proportional to the butanol concentration, as before [39]. 0.35 µm beads behaved identically to 0.5 µm beads, Fig. 1B (*Inset*). In stark contrast, the speed of 1.5 µm beads remained constant for up to 0.25% butanol, a concentration at which the speed of 0.5 µm and 0.35 µm beads had already decreased by ∼ 60%, Fig. 1B. Comparing the results of five bead sizes, we found that the effect of butanol on motor speed gradually decreased as the load increased, starting from 0.75 µm beads, Fig. 1C. These results confirm that the proportionality between PMF and motor speed does not hold across all viscous loads.

To exclude the effect of cell-to-cell variability in PMF, or in response to butanol, we simultaneously measured the speeds of two motors located on the same cell but operating under different viscous loads, similar to one of the previous experiments that demonstrated the linear relationship between motor speed and PMF [29], Fig. 2A. To do so, we cultured cells in the presence of sub-inhibitory concentrations of cephalexin, an antibiotic that induces filamentation in bacteria [40] (*Materials and Methods*). We then attached a 0.5 µm bead to one motor, and a 1.5 µm bead to a different motor on the same bacterium, which was sufficiently elongated to ensure the absence of direct contact between the two beads, SI Video 1. We treated those cells with a range of butanol concentrations (0-1%) in MM9+0.3% glucose, and found, consistent with our previous observations, that motor speeds were comparatively less sensitive to butanol treatment under high load, SI Videos 1 & 2. In particular, plotting the highload (1.5 µm) speed as a function of the low-load (0.5 µm) speed for motors on the same cells clearly showed that the high-load speed saturates at higher PMF, Fig. 2B, Fig. SI 3.

**Fig. 2.**
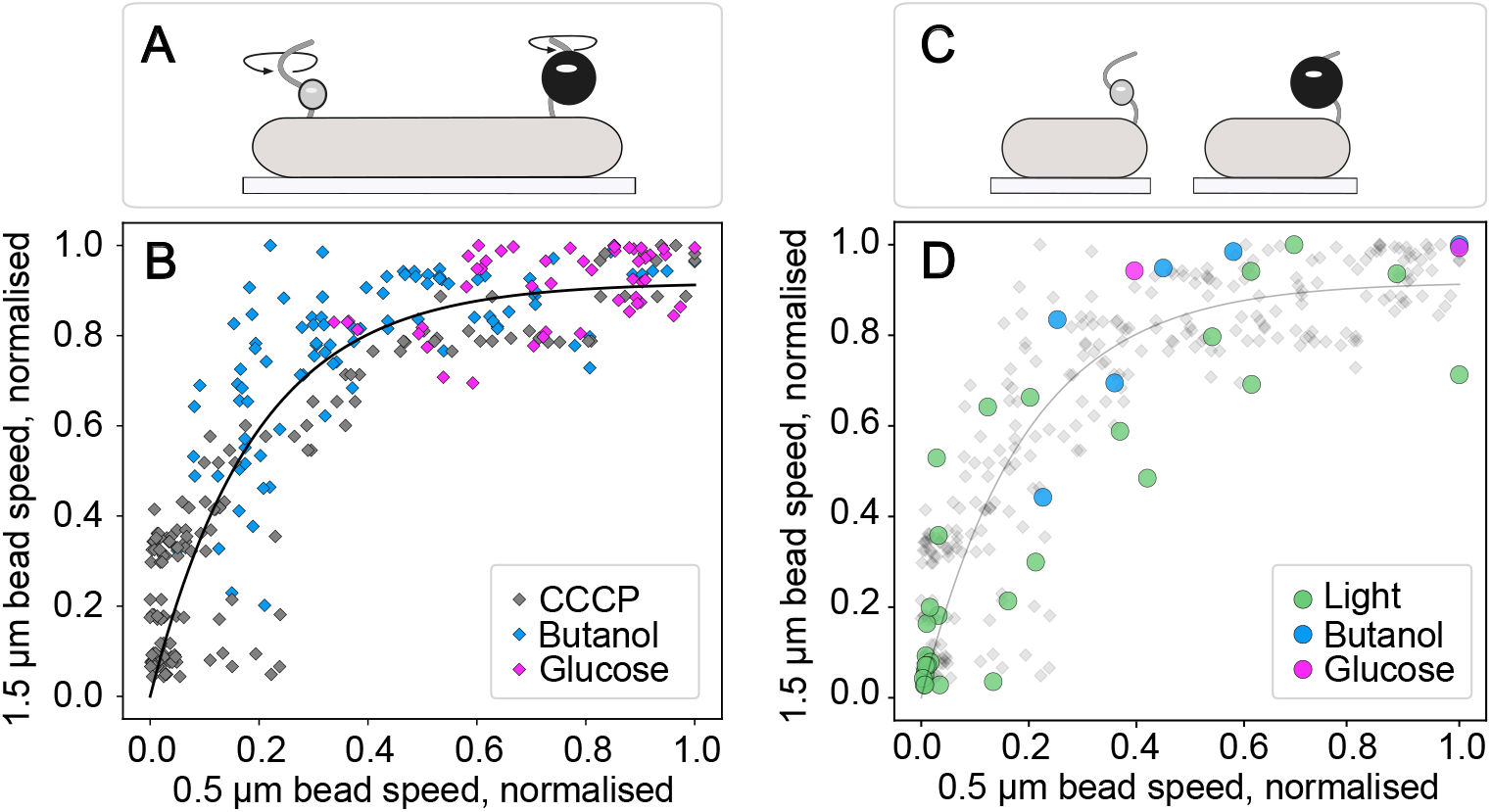
Speeds of two motors operating under different loads are not linearly proportional. **A**. A cartoon illustrating the one cell-two motors setup. Filamentous *E. coli* cell, grown in presence of sub-inhibitory concentrations of cephalexin, has two beads attached to two individual active motors, SI Videos 1&2. The speeds of both motors are recorded simultaneously while the cell is being exposed to PMF-altering treatments. **B**. Normalised speeds of two motors belonging to one cell as illustrated in A, respectively loaded with a 0.5 µm and 1.5 µm bead, and treated with CCCP (grey), butanol (blue), or glucose (pink) in order to manipulate the magnitude of the PMF. Black line shows an exponential fit applied to all the data to illustrate the saturation and facilitate comparison between data sets 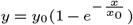, where *y*_0_ = 0.917 and *x*_0_ = 0.1925 are fitted parameters. **C**. A cartoon illustrating the single cell-single motor setup. *E. coli* cells are grown and prepared normally (see *Materials and Methods*), with either 0.5 µm or 1.5 µm beads attached to the truncated flagella filaments. Speeds of multiple motors are recorded independently. **D**. Averages of single cell-single motor data collected for motors loaded with either 0.5 µm or 1.5 µm beads, as shown in C, for a given butanol (blue) or glucose (pink) concentration from the data sets in Fig. 1. Green shows the data from the cells exposed to photodamage by 395 nm light at different powers given in Fig. SI 4C and D. In grey, in the background, single filamentous cell data from panel B as well as their exponential fit are given for reference. For **B**. and **D**. see *Materials and Methods* for description of the datasets and normalisation procedures, and Tables SI 1 - SI 2 for sample sizes.

Next, we further confirmed that the observed saturation is independent of the means by which we alter the PMF. While still working with individual filamentous cells, each with two motors loaded with differently sized beads, we decreased the PMF by exposing fully energised cells to the commonly employed protonophore Carbonyl Cyanide m-Chlorophenylhydrazone (CCCP) [29, 41], or increased it by supplying glucose to starved cells. In both cases, we recorded the speed of two motors under different loads as a function of time (Fig. SI 4A and B), and found that the data fall onto a single master curve, Fig. 2B. When we recorded the speeds of two 0.5 µm beads attached to a single cell, i.e. motors operating under the same load and PMF, we found that their speeds always varied linearly with each other, Fig. SI 5.

As a final method of altering PMF, we employed light of short wavelength (395 nm) to decrease the PMF via photodamage [39]. We used normal, non-filamentous cells, with either one small or one large bead attached to one of their motors, Fig. 2C. Once again, we found that the motor speeds decay was slower at higher loads than at lower loads (Fig. SI 4C and D). In Fig. 2D we show that all our results thus far: the speeds from elongated cells with two motors, or normal-sized cells with single motors, treated with all the different means of altering PMF, follow the same master curve.

Taken together, our observations suggest the PMF-speed proportionality is maintained across the physiological range of PMF values at lower, but saturates at higher loads. We sought to confirm this by directly measuring the PMF during glucose, butanol, and CCCP treatment. PMF is composed of the proton concentration gradient and the electrical potential across *E. coli*’s inner membrane [42]. Because *E. coli* maintains near neutral cytoplasmic pH and our experiments are performed in extracellular environment of pH=7, the dominant contribution to the PMF in our experiments comes from the membrane potential (*V*_m_). To measure it, we use PRoteorhodopsin Optical Proton Sensor (PROPS), and tetramethylrhodamine methyl ester (TMRM). The former is a genetically-encoded sensor previously validated in *E. coli* [43], whose relative fluorescence increases as the *V*_m_ becomes less negative. The latter is a fluorescent dye also previously validated in *E. coli* that behaves as a Nernstian sensor and gives quantitative measurement of *V*_m_ [33]. We were able to load TMRM only under the conditions described in [33], but not under our experimental conditions, Fig. SI 6. We attribute this to condition-dependent differences in cell membrane permeability or efflux activity. However, as expected, PROPS fluorescence increased in the presence of butanol or CCCP, and decreased upon glucose addition, see Supplementary Information (SI) text, Figs. SI 7, SI 8, & SI 9, and Table SI 3. An extended analysis of relative changes in *V*_m_, using also the PROPS calibration recorded in [43], supports our hypothesis that the motor speed remains proportional to PMF at low load, SI text and Fig SI 10.

### PMF-speed linearity is limited by the maximum torque

Since the motor speed saturation occurred only at high load, we hypothesised that the relationship between the PMF and the motor speed is limited by the maximum torque the BFM can produce, *T*_max_. To test this, in Fig. 3 we replot the motor speed data of Fig. 1C as torque-PMF curves (see *Materials and Methods* for torque calculation). We converted butanol concentrations to relative PMF values by assuming that the BFM speed is proportional to the PMF for sufficiently low loads, that is, 0.35 µm and 0.5 µm beads whose speed response to butanol overlaps, Fig. SI 11. We further assign a relative value of 1 to the maximum PMF of our experiments, which was obtained in MM9+0.3% glucose at 0% butanol and is thus presumably close to the maximum physiological PMF.

**Fig. 3.**
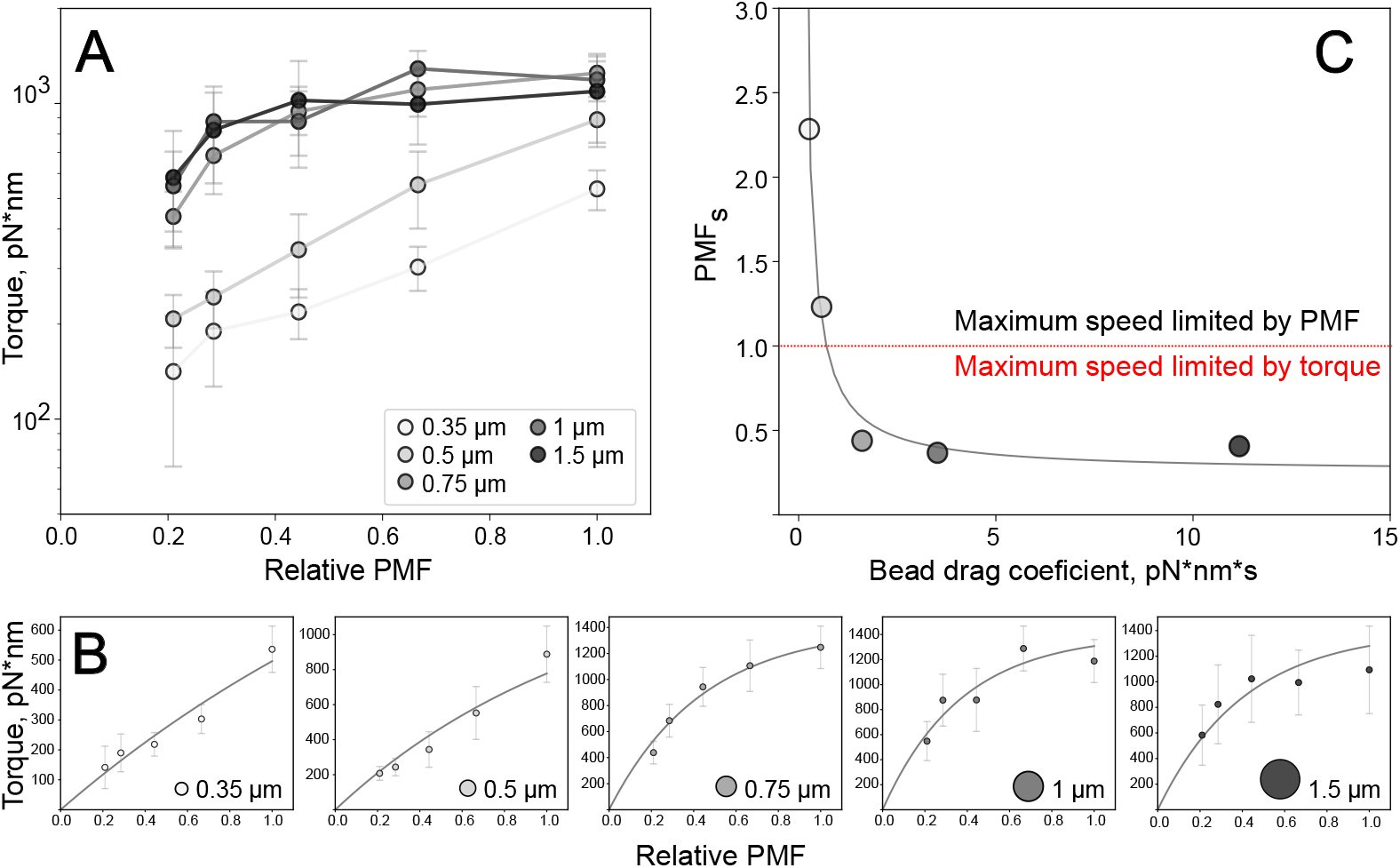
BFM torque saturates at high viscous load and high PMF. **A**. BFM torque is plotted against relative PMF. Torque and relative PMF are calculated as described in *Materials and Methods*. The colour scale for beads of different sizes is given in the inset. For larger beads and at higher PMF, the torque reaches the same saturating value of 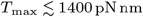 **B**. Individual Torque-PMF curves with exponential fits: 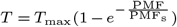, with *T*_max_ set to 1400 pN nm. Cartoons of different bead sizes given in the bottom right corners are to scale and the same colour scale as in (A) is used. **C**. PMF_s_ is plotted against the drag coefficient of the beads, *f*_*b*_, calculated as described in *Materials and Methods*. For loads with PMF_s_ >1 (0.35 µm and 0.5 µm beads), the maximum speed is not limited by torque for the whole range of physiological PMF values, whereas for loads with PMF_s_ <1 the maximum speed is torque-limited and saturates at high PMF. Error bars are standard deviations, and Table SI 1 gives sample sizes.

The torque-PMF curves for 0.75, 1, and 1.5 µm beads all converge to the same torque at high PMF, Fig. 3A. We fitted those curves with an exponential function,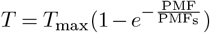 with 2 free parameters, *T*_max_ and PMF_s_, and found *T*_max_ for these three bead sizes to be 1430, 1313, and 1108 pN nm, respectively. In Fig. 3B we show the individual torque-PMF curves recorded for all bead sizes, where we now fit the same exponential function but set *T*_max_ to 1400 pN nm, keeping PMF_s_ as the only free parameter. For each viscous load, the fits returned PMF_s_, the relative PMF above which the torque-PMF relationship begins to saturate, Fig. 3C. For viscous loads characterised by PMF_s_>1, the saturation does not occur at physiological PMF and the BFM speed is proportional to PMF. For loads at which PMF_s_<1 the relationship is nonlinear; the motor speed is limited by the maximum torque and becomes less sensitive to PMF for PMF > PMF_s_, resulting in saturation and full insensitivity to PMF at higher PMF values.

Although we set *T*_max_ to 1400 pN nm in the fits of Fig. 3, we note that motor speeds and, thus, our calculated values of torque and, consequently fits of maximum torque, varied somewhat between experiments (all within previous estimates for the limiting torque of the BFM [12, 26, 34]). We discuss it in detail in SI text. Despite this variability in absolute numbers, the non-linearity of the PMF-speed relationship at high loads was consistently observed in all experiments. All our observations thus demonstrate that the saturation of the PMF-speed proportionality is defined by the maximum torque that the motor can produce.

### Fully-energised *E. coli* swim in the near-saturating regime

We next asked whether *E. coli* can experience this saturation while swimming in aqueous media. Since saturation occurs at high mechanical loads, the answer depends on the load experienced by the BFM of swimming cells, for which a wide range of assumptions and estimates exist in the literature. Several studies report that the motors of swimming *E. coli* operate within or close to their low-speed, constant torque plateau [44–46], while others propose that they operate at much lower torque [47–49]. The first situation would make the swimming speed PMF relationship prone to saturation, whereas in the second case the swimming speed should remain proportional to PMF. To obtain the answer, we measured the swimming speed of cells exposed to various butanol concentrations using differential dynamic microscopy (DDM) [37, 50, 51]. By comparing swimming speeds and motor speeds, each normalised by their respective value recorded in the absence of butanol, we found that the speed response curve of free-swimming cells lies between that of 0.5 µm and 0.75 µm beads, Fig. 4A. The butanol-speed curve for the free swimmers deviated slightly from the non-saturated 0.5 µm beads curve, indicating that the motors operate close to their saturating torque at maximum PMF. The addition of 8% Ficoll to the medium increased its viscosity ∼ 3.5fold (see *Materials and Methods &* SI text), consistent with previous measurements [25, 52]. The swimming speed response to butanol at this increased viscosity precisely matched the behaviour of 1 µm beads, Fig. 4B. Assuming the load with Ficoll to be equivalent to that of a 1 µm bead in Ficoll-free medium, and taking *η*_Ficoll_ = 3.5*η*, we found that the motors of cells swimming in Ficoll-free medium operate under a load equivalent to a 0.6 µm bead (see SI text), in line with the results of Fig. 4A. As in the double bead experiment of Fig. 2, high-load swimming speed in Ficoll exhibited saturation at high PMF compared to the low-load swimming speed, Fig. 4B (*Inset*).

**Fig. 4.**
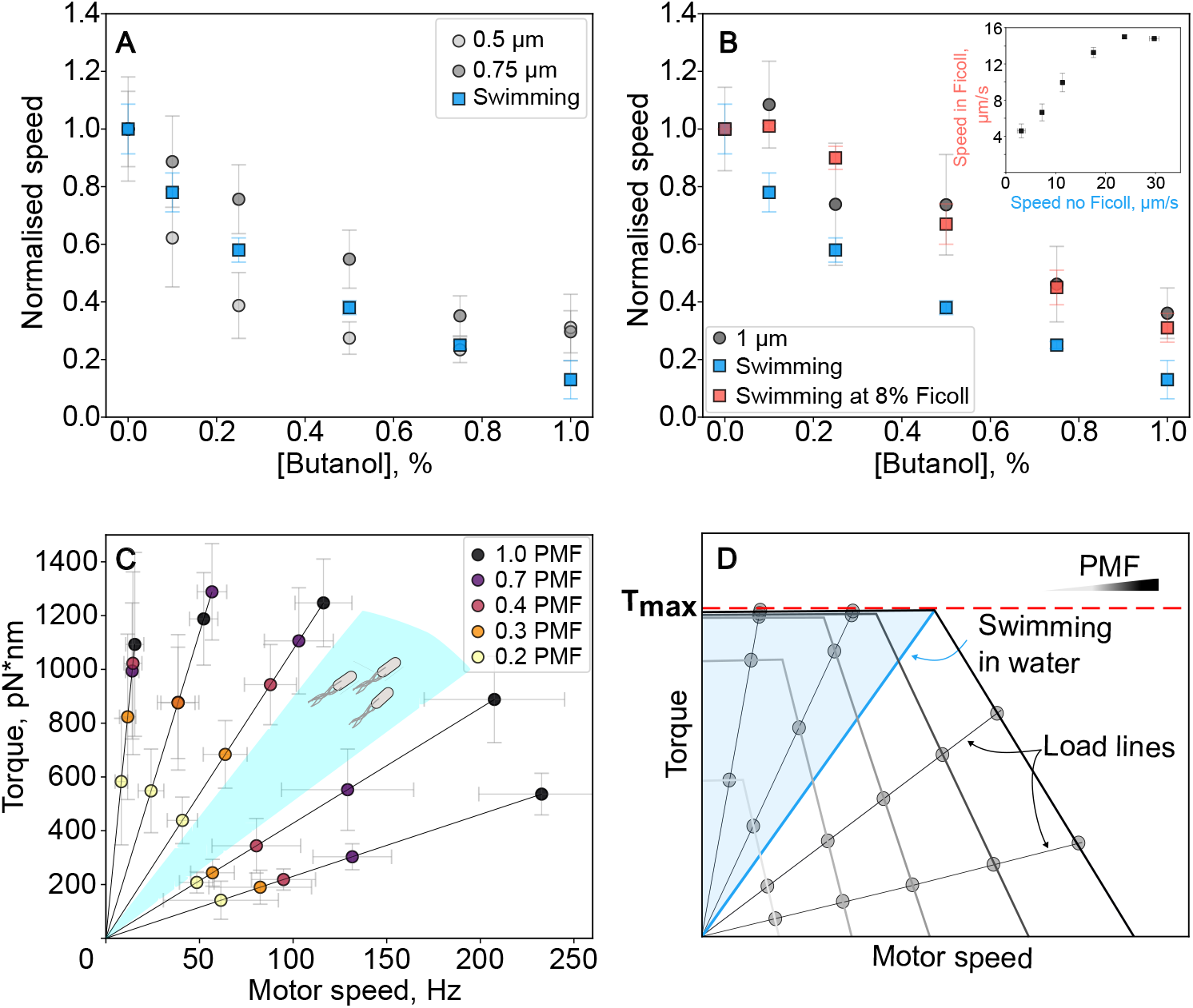
Free swimming cells produce near-saturation torque. **A**. The swimming speed response to increasing butanol concentrations is equivalent to the response of BFM under the viscous load imposed by ∼ a 0.6 µm in diameter bead. **B**. Addition of 8% Ficoll increases viscosity 3.5-fold, bringing the load on the motor to that imposed by a 1 µm bead. (***Inset***) Swimming speeds of cells swimming with and without Ficoll are not linearly proportional at high PMF. **C**. Torque-speed curves of the BFM at different PMF values. Black lines represent load lines for beads of different sizes. Cyan shaded region is the torque-speed space occupied by bacteria swimming freely in an aqueous environment with a viscosity similar to that of water. **D**. A revised cartoon of the idealised torque-speed curves, each depicting BFM torque-speed characteristics at different PMF values (given by the grey colour scale in the top right corner). Straight lines represent load lines. The load line that corresponds to swimming in water shows the lower boundary of the swimming regime, and the light blue shaded region shows the torque-speed space available to *E. coli* when swimming at higher viscosities. Error bars are standard deviations. See *Materials and Methods* normalisation procedures, and Table SI 1 for sample sizes.

Consistent with our hypothesis that the PMF-speed proportionality breaks at higher PMF values but holds at lower PMF (below 45% of PMF_max_), normalising the swimming speeds obtained with and without Ficoll by their respective values at 0.25% butanol, equivalent to 44% PMF_max_, collapses the speed-butanol curves onto a single curve at low PMF, Fig. SI 12.

The motor torque-speed curves obtained in the butanol experiments allowed us to place swimming cells (in the absence of Ficoll) onto the torque-speed plane, Fig. 4C. Swimming in water (or other aqueous environments like MM9) occurs at torques below or close to the “knee” of the torque-speed curves, and approaches the limiting torque at PMF_max_. Exponential fits of these experimental torque-speed curves are presented in Fig. SI 13.

Taken together, our results revise the well-known idealized torque-speed diagram, where the separation between individual torque-speed curves is proportional to the difference in PMF between them [27, 29]. The updated diagram includes the maximum torque, Fig. 4D, and the torque-speed curves in low-speed, high-torque regimes now collapse on top of each other. The lower limit of the viscous load experienced by free-swimming cells is estimated for an aqueous environment of viscosity similar to that of water. At higher viscosities, the swimming load line shifts to the blue-shaded region in Fig. 4D.

## Discussion

The bacterial flagellar motor is a sophisticated molecular machine whose structure and operational principles have been extensively studied [6]. However, discovery of new features of the motor forces us to revisit questions long believed to be answered. Here, we addressed the well-known PMF-speed proportionality and found that in *E. coli* it is limited by the maximum motor torque, and that this limit is reached within physiological values of the PMF. Although the existence of such maximum torque is not surprising, the fact that it can be reached already at 45% of maximum PMF, Fig. 3A, was unexpected. Consequently, in Fig. 4D we modified the previously reported idealised torque-speed curves [27, 29] to include our findings.

### Maximum torque in the context of previous work

Our results contradict previous measurements showing that the BFM speed and PMF are linearly proportional irrespective of the load on the motor [24, 29], for which we propose several explanations as follows.

We first tested the possibility that our results are specific to our strain or experimental conditions. Strain KAF95, used extensively in previous works [25,29,53,54], harbours a *cheY* deletion that causes its flagellar motors to rotate exclusively counter-clockwise (CCW), whereas strain EK01 used here has a wild type MG1655 motor capable of switching between clockwise (CW) and counter-clockwise rotation. In Fig. SI 14A and B, we show that the saturation at high load is observed in EK01 irrespectively of the direction of rotation. Additionally, we found that our observations are reproducible in strain KAF95 grown similarly to previous work (see *Materials and Methods*) [25, 29, 53, 54], Fig. SI 14C and D. Although the Fung and Berg publication used the HCB892 strain, with deleted F_1_F_o_-ATP synthase and grown in LB with fructose, which we were unable to obtain and therefore have not tested, overall we conclude that strain differences do not explain the observed discrepancy.

In Gabel and Berg’s experiment, the rotation of two motors was simultaneously observed in individual KAF95 cells [29]. One motor was anchored to the surface via the filament, in a so-called tethered cell configuration, driving the rotation of the entire cell body, which represents a large viscous load. The other motor was rotating under a significantly lighter viscous load imposed by a 0.4 µm bead. The cell was chemically treated to collapse the PMF and the rotations of both the cell body and the bead were detected, showing, in contrast with our results, proportionality across the range of PMF values tested. This discrepancy is not linked to different experimental time scales, because the characteristic time of the PMF change in [29] (∼ 5 − 10 min) is within the range covered in this work (∼ 2 − 30 min). This is about an order of magnitude more than the time required for stator unit association to the motor (20 s), see Table SI 4. Thus, both here and in [29], the total speed change corresponds to that of a motor in steady state and results from the combination of single stator PMF-torque response and stator dynamics.

The discrepancy with our observations can instead be ex-plained by the slow body rotation not being driven by an active motor tethered to the slide, but rather being a passive counter-rotation balancing the bead’s active rotation. In this configuration, the cell would be tethered either by a broken motor or by a hydrophobic patch allowing the cell to pivot passively, even in the absence of a flagellum [55]. The observed speeds and the ratio of cell-to-bead rotational friction coefficients are consistent with this explanation (see SI text). Repeating the experiment of [29] with beads of various sizes and a strain whose motors rotate exclusively counterclockwise (see *Materials and Methods*), we found that at least 25%, and up to ∼ 80% of rotating cells attached to a rotating bead were in fact passively counter-rotating, with significant variability between different preparations, Fig. SI 15 and Table SI 5. Passive counter-rotations were identified in slowed down movies from the fact that the cell body and the bead rotated in opposite directions, whereas they would rotate in the same direction if driven by two active motors, SI Videos 3 & 4. These passive counter-rotations were sta-ble for at least 15 min, SI Videos 5-8. A measurable passive counter-rotation is more likely the closer the cell and bead rotation axes are, which is the geometry enforced by the pinhole method of detection used in previous work [29]. This geometrical constraint also decreases the likelihood of simultaneously measuring two active motors: two of the cell’s ∼ 5 motors [49, 56] would need to be attached to either the glass slide or the bead, and would need to be aligned along the cell’s short axis. To confirm that this passive counter-rotation likely explains the results of [29], we collapsed the PMF by exposing cells to CCCP as in the previous study, and plotted the body frequency against the bead frequency, Fig.SI 16 and Table SI 5. As expected, passively counter-rotating cells produced a linear relationship similar to [29], whereas an actively rotating cell produced a non-linear relationship matching that of Fig. 2.

In Fung and Berg’s experiment [24], a cephalexin-treated filamentous cell was introduced into a micropipette, and the rotation of the motor located on the portion of the cell outside of the pipette was detected. The motor was operating under a high viscous load imposed by a non-filamentous cell attached to the motor, and a command voltage, calculated to result in 0 to −150 mV experienced by the motor, was applied with a microelectrode located inside the pipette to the filamentous cell made permeable by gramicidin S [24]. However, a possible additional voltage drop across the outer membrane (see SI text) and the conductivity of the periplasm were not considered, and strong indications that the voltage experienced by the motor was significantly lower than calculated can be found in David C. Fung’s and Christopher V. Gabel’s theses [57, 58]. In the former, the author states that the maximum motor speed achieved by powering bacteria with an external source is similar to the motor’s initial speed in a motility buffer with no carbon source [57]. Whereas the speed of free swimming cells increases by ∼ 60% after addition of 100 µM-50 mM glucose to motility buffer [37], indicating that the PMF in carbon-free motility buffer cannot be higher than 62.5% of the physiological maximum. In the latter, the author describes the attempts to repeat Fung’s experiments using a µm bead as a marker. The bead reached the maximum speed of 37 Hz, after which dielectric breakdown of the cell membrane occurred. However, under physiological PMF in the presence of lactic acid as carbon source, the author observed the 0.4 µm bead at speeds of 220 Hz [29]. Thus, the experimental evidence suggests that externally generated PMF in the micropipette experiments is significantly lower than the maximum physiological PMF of *E. coli*.

Apart from the two publications discussed above, direct or indirect evidence of non-linear PMF-speed relationship has been reported in several previous publications, and not always interpreted as such. Shioi *et al*. [59] and Khan *et al*. [60–62] reported that the relationship between swimming speed and PMF for *Bacillus subtilis* and *Streptococcus* exhibits a threshold at low PMF and a saturation at high PMF. The saturating PMF values reported by these publications (at high loads) are at ∼ −50 mV, whereas our measurement place it closer to ∼ −100 mV. Sowa *et al*. reported different rela-tive speed changes for Na^+^-driven *Vibrio alginolyticus* motor under low and high loads, with 0.6 µm and 1.08 µm beads respectively [63]. More recent work by Inoue *et al*. found the torque of the chimeric Na^+^-driven *E. coli* motor changes significantly with temperature at low load (0.35 µm beads), whereas it was constant at 2000 pN nm irrespective of temperature under high load (1 µm beads) [64]. Gupta *et al*. reported changes in the swimming speed of *E. coli* with addition of an ionophore, indole, that could not be reproduced in a single motor experiment with 2 µm latex beads [65]. Although alternative explanations were proposed in both studies, the results are also consistent with our finding that the BFM at high loads reaches its saturating torque. In fact, the PMF-speed relationship in Gupta *et al*. [65] obeys the same saturation law observed for high-load beads in Fig. 3B, see Fig. SI 17. And the results of Inoue *et al*. lead us to suggest that the chimeric, sodium-driven *E. coli* motor experiences a similar kind of torque-IMF nonlinearity. Overall, our results fit well in the context of existing literature, with plausible explanations for the seeming discrepancies.

### Implications for the models of the BFM function

The nonlinear nature of the PMF-speed dependency challenges current models of BFM function. Firstly, they all assume tight coupling between motor rotation and the PMF across the physiological range [7, 27, 66], which will now need to be reexamined. Secondly, we note that stator units are torque sensors [13, 14, 67]. When the speed increases due to an increase in the PMF, so does the torque, and additional stator units are recruited. The linear speed-PMF relationship of the whole motor at low torque is a result of the stator dynamics, and it is remarkable that the motor can achieve it, given its dynamic nature. Future theoretical explanations of how it does so will need to also include the new critical constraint on the whole-motor PMF-speed relationship set by our results. For example, mechanical limits of the rotor stator-complex may be considered, whereby the stator-rotor interface can only sustain some maximum force before mechanical slipping. Similarly, there could be a limit to conformational transitions, e.g. causing a finite torque per stator.

Our estimates of maximum torque from all our data, 1108-1839 pN nm (see SI text), are within the range of previously published values for a wild type *E. coli* motor: Reid *et al*. estimated the maximum torque to be 1260 *±* 190 pN nm [12], Ryu *et. al* measured torque up to ∼ 1300 pN nm [26], and Berry *et al*. reported a stall torque of 4500 pN nm [68], which was later corrected to ∼ 2000 pN nm [6, 34]. Estimates of the torque produced by a single stator unit: 146 [12], 250 [69], or 320 pN nm [26], with up to 11-12 stator units per motor [12], are also in line with our results. Here, for better comparison with previous torque estimates, we applied the equation most commonly used for torque calculation in the bead assay (see *Materials and Methods*) [15, 26, 28, 64], and did not include hydrodynamic surface effects between the cell body and the rotating bead [69]. We discuss in SI text the possible origins of the relatively large variability in our assay, but stress that our two main conclusions are robust to those variations: the saturation of the PMF-speed relationship at high loads was observed across all experiments, and the value of the swimming load (0.6 µm bead equivalent) obtained from butanol experiments was independent of the initial motor speed.

### Implications for BFM use as a physiological sensor

Our findings set new limitations on the use of the BFM as a physiological indicator of PMF [39, 43, 65, 70]. To use the PMF-speed proportionality, it is necessary to confirm that the motor operates below its limiting torque, the lower boundary of which we estimate as ∼ 1100 pN nm. As an additional test, we suggest the use of butanol or another ionophore (note that the commonly used protonophore CCCP should be used with care in *E. coli* as it is a substrate for efflux pumps [41]) to probe the sensitivity of motor speed to PMF variations at a given load and in a given condition. This considerationis especially relevant for the use of tethered cells as PMF probes. Their load is typically high but depends on the size of the cell and the geometry of attachment, which can lead to inconsistent results if the saturated regime is reached at different PMFs for different cells. With these limitations in mind, we remain convinced that the BFM in its linear PMF-speed regime is the best PMF sensor currently available [42].

In this manuscript, we chose to work with relative rather than absolute PMF variations, assuming that PMF in MM9+0.3% glucose is close to the maximum physiological value. While absolute PMF measurements in live Gram-negative bacteria such as *E. coli* remain challenging [42], and we have not been able to load the TMRM dye in our conditions, Fig. SI 6, some estimates of the PMF achieved by respiring *E. coli* are available: a theoretical study predicted a range of −170 mV to −229 mV from the free energies of NADH oxidation and ATP hydrolysis [71], and values of −230 mV and −160 mV were estimated from proton and potassium fluxes in *E. coli* spheroplasts [35], and by radioactive probing [36], respectively. Using those estimates, relative PMF variations recorded by BFM speed measurements can be converted to absolute PMF values.

### Implications for understanding how E. coli swims and why it has evolved to swim this way

All current studies attempting to understand bacterial swimming consider the torque produced by the flagellar motors to drive flagella, and balance it by the torque driving the rotation of the cell body, toensure zero net torque on the swimming bacterium. The torque produced by the motors depends on the load under which they operate. Theoretically predicting it is difficult, because load sharing between motors and dissipation by filament-body and filament-filament interactions must be taken into account [44, 72]. One way to measure this load is to measure the motor speed using bead assay [25, 27], and compare it to the rotation speed of the flagella bundle of swimming *E. coli* [44, 73]. However, we were unable to find both measurements done as part of one study. As a result, motor speeds have often been compared to flagella bundle speeds measured in different conditions or with a different strain. This explains the wide range of estimates for the load experienced by the motors of swimming cells currently used in the literature, with some studies assuming that all motors operate within the high-torque plateau [44, 45, 72], whereas others consider or report intermediate [48] or small loads [49]. Our direct comparison of the bead assay to swimming speed results enabled us to estimate this load precisely, the lowest being the equivalent of a 0.6 µm bead in the aqueous medium, see Fig. 4 and SI text for details. Because our butanol protocols offer a way to measure this load in a direct comparison with the fixed known load in a range of conditions, they should be of interest to the microbial swimming community.

Although our work shows that the PMF-speed saturation occurs at physiological PMF under medium-to-high load, it does not address why evolution sets the limiting torque to its current value. One possible explanation is that it is sufficiently high to never hamper *E. coli*’s chemotactic efficiency in natural environments. Indeed, while *E. coli*’s primary habitat, the gastrointestinal tract of warm-blooded animals, is presumably much more viscous than water, it is also mostly anaerobic, which means that *E. coli*’s PMF in this environment is likely significantly lower than in our aerobic experiments. At the same time, *E. coli* cells swimming outside a host presumably experience low viscous loads and are unlikely to be as fully energised as in our glucose experiments. In both cases, it is possible that a higher limiting torque would not significantly benefit the cells since a PMF high enough for saturation is never reached. Alternatively, maintaining a constant, PMF-independent speed in the saturated regime might confer advantages by preventing cells from accumulating in low-PMF regions in the absence of an attractant/repellent gradient, or by facilitating the collective migration of bacteria all swimming at the same speed, regardless of whether a cell is at the front of the band and has access to fresh nutrients, or is (transiently) behind and relies on endogenous reserves to generate a reduced PMF.

## Materials and Methods

### Bacteria culturing and media

#### Strains and media for BFM speed measurements

All strains and media used in this work are listed in Tables SI 6 and SI 7. We use the strain EK01, a derivative of MG1655 with the flagellin encoding *fliC* gene replaced by *fliC*^*sticky*^ [74] for most of the BFM speed experiments. The mutation makes flagella hydrophobic and allows polystyrene beads to stick to them by electrostatic interaction. For experiments with CCCP, we used EK01 Δ*tolC* [41]. For experiments validating the possible passive counter rotation of the cell body under the action of a rotating bead, we used EK01 Δ*cheY*. This deletion makes the motor rotate exclusively counterclockwise. For photodamage experiments, we use strain EK07, a derivative of EK01 with a chromosomally integrated pHluorin [39, 75]. The KAF95 strain was also used for comparison with previous work [29]. This RP437 derivative strain bears a *fliC* and *cheY* deletion, as well as an ampicillin-resistant plasmid encoding the *fliC*^*sticky*^ mutation.

For all experiments apart from those using KAF95 strain or replicating the geometry of [29], bacteria were grown in Lysogeny Broth (Table SI 7). An aliquot of overnight culture (OD=5.5) frozen and stored at −80°C in the presence of 20% glycerol was thawed and diluted in fresh LB to OD ≈ 0.003 (1:1000 dilution of the overnight culture) and grown at 37°C with shaking (220 rpm) to OD = 2.0. After harvesting, cells were “sheared” by multiple passing through a narrow gauge needle [76], and washed 3 times by centrifugation (8000 g, 2 min) into Modified M9 (MM9, Table SI 7), pH=7 for all apart from photodamage experiments where pH=7.5, with or without glucose (0.3%).

To measure the speed of multiple motors on elongated cells, cells of EK01 (butanol and glucose experiments) or EK01 Δ*tolC* (CCCP experiments) were grown as above, except that 50 µL of a 10 mgmL^−1^ cephalexin aqueous solution was added to the growth medium (50 µgmL^−1^ final cephalexin concentration) when the culture reached OD ≈ 1. Cells were grown for an additional 45-60 min to OD ≈ 2 before being harvested and prepared for the bead assay in MM9, with or without glucose (0.3%), supplemented with 50 µgmL^−1^ cephalexin.

Strain KAF95 was grown in Tryptone Broth (TB, Table SI 7.). A frozen stock was thawed and diluted in fresh TB to OD ≈ 0.003 (1:1000 dilution) and grown at 30°C with shaking (220 rpm) to OD ≈ 0.4. After harvesting, cells were “sheared” as described above and washed 3 times by centrifugation in Motility Buffer (MB, Table SI 7), pH=7, with or without glucose (0.3%).

For experiments replicating the geometry of reference [29], strain EK01 Δ*cheY* was inoculated from frozen stock in LB and grown overnight at 37°C with shaking at 220 rpm. The overnight culture was diluted in TB to OD = 0.04 (∼ 1:200 dilution) and grown at 30°C with shaking (220 rpm) to OD ≈ 0.4. Cells were “sheared” as described above and washed 3 times by centrifugation in MB or in Gabel’s Motility Buffer [29] (GMB, Table SI 7), pH=7, with or without glucose (0.3%).

#### Strains and media for membrane potential measurements

We used strain YD133, the Keio Collection BW25113 derivative with Δ*fliC*, Δ*pilA*, Δ*flgE* [77], transformed with plasmid pWR20 (GFP expression) [78] and an arabinose-inducible pBAD plasmid for PROPS expression [43] (Table SI 6). For TMRM experiments, we used EK01 Δ*tolC*. For PROPS experiments, a pre-culture was grown from a frozen stock at 1:1000 dilution in LB supplemented with 100 µgmL^−1^ ampicillin at 37°C with shaking (220 rpm). The overnight culture was diluted 1:1000 in fresh LB supplemented with 100 µgmL^−1^ ampicillin, 5 µM all-trans retinal and 0.0005% by weight arabinose, and grown at 37°C with shaking (220 rpm) for 5h 30 min to OD=2.3. After harvesting, cells were washed 3 times by centrifugation (8000 g, 2 min) into MM9 supplemented with 5 µM all-trans retinal, with or without glucose. Cells were stored on ice until the experiment. All media used in PROPS experiments contained 5 µM all-trans retinal [43].

For TMRM experiments, a first culture of EK01 Δ*tolC* (Culture 1) was grown in LB to OD=2.2 at 37 ^◦^C, following the protocol used for the bead assay. A second culture (Culture 2) was prepared using a protocol similar to that used in a previous TMRM study [33]. Briefly, 60 µL of a frozen stock of EK01 Δ*tolC* was inoculated in 10 mL of TB and grown for 5.5 h at 30 ^◦^C to OD=0.24. Cells of Culture 1 were washed twice by centrifugation (8000 g, 2 min) into MM9 devoid of Ca^2+^ and Mg^2+^ ions. Cells of Culture 2 were washed twice in MB, pH=7-7.2. To increase the cells’ permeability to TMRM [33], cells of Culture 1 were washed by centrifugation a third time in MM9 lacking Ca^2+^ and Mg^2+^ and supplemented with 10 mM EDTA, pH=7-7.2. Cells of Culture 2 were washed in MB supplemented with 10 mM EDTA, pH=7.2. Cells were incubated for 10 min at room temperature in the EDTA-supplemented media, before being washed three times in either MM9 lacking Ca^2+^ and Mg^2+^ (Culture 1) or MB (Culture 2). Both suspensions were diluted to OD=0.1 in their respective buffers, and were supplemented with 100 nM TMRM by serial dilution of a 50 mM TMRM stock solution in DMSO. Cells were incubated with TMRM for 30 min at room temperature before being introduced into tunnel slides for fluorescence imaging.

#### Swimming speed measurements

We conducted them with strain AD80, a smooth-swimming MG1655 mutant constructed by deleting the *cheY* gene [37] (Table SI 6). The bacteria were grown as for motor speed measurements, to OD = 2 before being harvested and washed by filtration to preserve the filaments [37] in MM9+0.3% glucose. The concentrated suspension obtained was diluted with MM9+0.3% glucose and stored on ice until differential dynamic microscopy measurement.

### Data acquisition

#### BFM speed recording with bead assay

For the BFM speed recordings, we used the bead assay [21] in combination with back-focal-plane interferometry (BFP) [3, 79, 80] or a highspeed camera (iDS uEye CP, Teledyne Photometrics Prime 95B and Andor Zyla DG-152V-C1E-FI) for all bead sizes. BFP interferometry has a higher temporal resolution but a lower throughput than the camera. Both methods were used to ensure they returned identical results, but most of the data were collected with the cameras, and the data sets were later merged into a single speed distribution for each experimental condition, Fig. SI 1 & SI 2.

For the bead assay, cells were immobilised on glass tunnel slides coated with poly-L-lysine (PLL) or, for photodamage experiments, a custom-built flow cell, both prepared as described previously [3, 39, 81]. The exception are experiments replicating the geometry of [29], where no PLL was used. Polystyrene beads of defined sizes were then attached to their truncated flagella [81], see the actual bead sizes in the Table SI 8. In the tunnel slides, the medium was refreshed every 5-10 min by flushing ∼ 50 - 100 µL of fresh medium to ensure that glucose and oxygen were not depleted during recording. For photodamage experiments, the flow cell was constantly supplemented with fresh medium using a peristaltic pump (Fusion400, Chemyx, USA) at a flow rate of 10 µl/min.

For single-motor, single-cell bead assay where we used a high-speed camera, the frame rate was 460 fps for 1.0 µm and 1.5 µm beads, or 1200 fps for all bead sizes (1.0 µm and 1.5 µm beads were recorded with both settings). For the bead assay with cephalexin-treated cells, the frame rate was 1200 fps. In experiments replicating the geometry of [29], the beads’ and cells’ trajectories were recorded at 1230 fps for 0.5 µm beads, 500 fps and 992 fps for 0.75 µm beads, 100 fps and 200 fps for 1.5 µm beads.

For butanol experiments, cells immobilised in tunnel slides were exposed to MM9+0.3% glucose supplemented with appropriate concentrations of 1-butanol (molecular biology grade, Sigma-Aldrich), and 10 s-long videos were recorded. For *in-situ* glucose treatment of cephalexin-treated cells, cells were prepared and immobilised in MM9. MM9+0.3% glucose was flushed in the tunnel slide a *t* = 0 min, and 40 s videos were taken every 1 min for the first 15 min, then every 5 min. For CCCP treatment of cephalexin-treated cells, a continuous 150 s video was recorded and 50-100 µL of concentrated CCCP solution (10-50 µM) in MM9+0.3% glucose was deposited on one side of the chamber at *t* ∼ 10 - 15 s, allowing CCCP to diffuse in and progressively dissipate the cells’ PMF. For CCCP treatment in experiments replicating the geometry of [29], the same procedure was followed using 50 µL of concentrated CCCP solution (25 µM) in MB+0.3% glucose or GMB+0.3% glucose, and 200 s videos were recorded. To determine the fraction of active/passive cell body rotations in the experiments replicating the geometry of [29], 10 s videos were recorded.

For BFP interferometry, the time course of the bead rotation was recorded with a position-sensitive detector (PSD Model 2931, New Focus, USA) at 10 kHz sampling rate for 10 s per motor for all experiments, except those involving light damage. For the latter, the motor speed was recorded for an initial 30 s period without light exposure followed by continuous exposure to 395 nm with a narrow-spectrum UV LED (Cairn Research, Faversham, UK). Once the light was activated, the recording continued for 15 min.

#### Differential dynamic microscopy

For swimming speed mea-surements in the absence of Ficoll, washed cells were stored at OD = 1 on ice. For each butanol concentration, 150 µL of this suspension was diluted to OD = 0.3 with 350 µL of MM9+0.3% glucose supplemented with an appropriate butanol concentration. 200 µL of this suspension was used immediately to fill a rectangle glass capillary (VitroCom, inner diameters 0.4 mm *×* 8 mm). The capillary was sealed with petroleum jelly (Vaseline) to prevent evaporation, and bacteria were imaged with a phase contrast objective (Plan Fluor 10×/0.3), approximately 150 µm above the bottom of the capillaries. 40-s long movies were recorded every 5-6 min at 100 fps using an Orca Flash 4.0 camera (Hamamatsu) mounted on a Nikon Ti-E microscope.

For experiments with Ficoll 400 (Sigma Aldrich), washed cells were stored at OD = 3 on ice. 50 µL of this suspension was diluted to OD = 0.3 with 450 µL MM9+0.3% glucose supplemented with an appropriate butanol concentration and 9 wt% Ficoll (resulting in 8.1 wt% final Ficoll concentration). Capillaries were prepared and imaged as described above.

For viscosity measurements, 0.5 µm and 1 µm polystyrene beads (2.5 wt% initial concentration) were diluted 10x in either deionized water or MM9+0.3% glucose +9 wt% Ficoll (8 wt% final Ficoll concentration), and the resulting suspensions were imaged using the same conditions as for the swimming speed measurements.

#### Membrane potential measurements

Cells harbouring PROPS were immobilised on PLL-coated glass tunnel slides, and suitable regions with cells lying flat on the glass substrate were selected by visual inspection of phase contrast images. For each measurement, four images were taken with the Teledyne Photometrics Prime 95B camera, using a phase contrast objective (Plan Apo DM Lambda 100x/NA=1.45), as follows: one phase contrast image, one GFP image, and two images for PROPS fluorescence. We used the green channel of a pE-300 LED (CoolLED, UK) at 100% power for PROPS, the blue channel at 5% power for GFP, and appropriate emission/excitation filters in filter cubes. For PROPS, these were an ET545/30x filter (Chroma, USA) for excitation and LF635-B filter (Semrock, USA) for emission. A GFP filter set (Excitation: ET470/40x, Chroma. Emission: 500/LP, Semrock Brightline) was used to image GFP. PROPS images were collected with 100 ms and 1 s exposure times. GFP images were collected with 5 ms exposure time.

To evaluate the effect of butanol on PROPS fluorescence, cells were suspended in MM9+0.3% glucose and exposed to various concentrations of butanol in a tunnel slide (0%-2%). The effect of CCCP was observed similarly by exposing cells in the tunnel slide to MM9 supplemented with a CCCP concentration large enough to fully dissipate PMF (≈ 250 µM). Finally, for glucose measurements we first measured the cells’ fluorescence in the absence of glucose, added the carbon source, and then measured the same cells 3 times over a period of 15 min, every 5 min. The first point was measured 5 min after glucose addition.

Data were collected on two consecutive days using the same batch of cells. On the first day, this was the fresh batch, which was stored overnight at 4°C for the second day. PROPS is stable for at least a week in *E. coli* [43], and no differences were observed between the two data sets.

For TMRM measurements, cells incubated for 30 min in TMRM-supplemented media were immobilised on PLL-coated glass coverslips, either in MM9 + 100 nM TMRM lacking Ca^2+^ and Mg^2+^ (Culture 1), or in MB + 100 nM TMRM (Culture 2). Images were recorded using the same setup as for PROPS measurements, with the LED at 10% power, a DsRed filter set (Excitation: ET545/30x, Chroma. Emission: ET620/60m, Chroma), and 50 ms exposure time. For both cultures, a first dataset was recorded immediately after cells were introduced into the tunnel slide. A second dataset was recorded 10 min later, immediately after washing unattached cells out of the tunnel slide with either MM9 + 100 nM TMRM lacking Ca^2+^ and Mg^2+^ (Culture 1), or MB + 100 nM TMRM (Culture 2).

#### Effective power calculations

To measure the total effective power delivered to the cells during light exposure, a power meter with a photodiode sensor (PM100D and S120C, respectively, both from Thorlabs, UK) were used to quantify the illumination power (mW) emitted by the LED at the sample plane, and the total power was calculated as described in [39].

### Data analysis

#### Single motor’s speed calculation

BFP interferometry output was given in x-y coordinates calculated from the PSD photocurrents as described in [39]. For high-speed camera recordings, a region of interest (ROI) was manually selected in FIJI/ImageJ, Otsu threshold was applied to each ROI, and the coordinates of the center of mass of the binary image were calculated for each frame giving the x-y time series of the bead rotation. Alternatively, bead trajectories were recorded via direct tracking in MATLAB R2022a, using a code adapted from [82]. Both methods returned identical results when applied to the same data sets.

The rotational frequencies were then extracted by using a fast Fourier transform (FFT) on the x-y series. A flat-top window discrete Fourier transform (window size of 16384 data points for BFP interferometry data and 1024 points for high-speed camera data, 0.01 s moving window steps) was applied to the bead coordinates to obtain a time series of motor speeds. The mean speed was calculated for each motor, SI 1 and SI 2.

For experiments with EK01 Δ*cheY* replicating the geometry of [29], if both the cell body and bead rotated clockwise in the field of view (corresponding to counter-clockwise rotation of the motors), these were deemed two actively rotating motors. CW bead rotation and CCW body rotation was taken as passive body counter-rotation (see SI Text). In a few cases, bead rotation was undetermined, for example when a bead located on the side of the cell body produced a projected trajectory that appeared CCW. The cell’s rotation caused the bead to go in and out of focus, precluding automated analysis. Thus, bead and cell trajectories were analysed manually. At multiple time points in acquired movies, the cell’s rotation speed was measured from the number of frames needed for the cell to rotate by 90°, and the bead’s rotation speed was calculated from the number of rotations performed during this time interval.

#### Torque calculations

The torque was calculated as before [15, 26, 28, 64]:

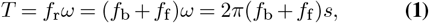

where *T* is torque, *f*_r_ is the rotational frictional drag coefficient and *ω* is the angular velocity of the BFM, i.e. its speed (*s*) obtained by FFT in Hz and multiplied by 2*π. f*_r_ is the sum of the drag coefficients of the flagellar filament stub (*f*_f_) and the attached bead (*f*_b_). The filament stub contribution was estimated as *f*_f_ ≈ 0.1 pN nm Hz^−1^ (mean of values reported in [64] and [83] divided by 2*π*). *f*_b_ was calculated as:

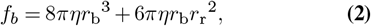

where *r*_b_ is the radius of the bead, *r*_r_ is the eccentricity of the bead’s trajectory and *η* is the viscosity. *η* was set to 1 mPas and *r*_r_ to 200 nm (values measured from the videos were 100-320 nm).

Note that multiple studies used frequency instead of angular velocity in Eq. 1, thus, having to multiply the rotational frictional drag coefficient by 2*π*.

#### Swimming speed and viscosity measurements

DDM movies were analysed using previously described methods [50, 51]. The average speed of the suspension was obtained for each movie by assuming that the speed distribution of motile *E. coli* cells takes Schulz form:

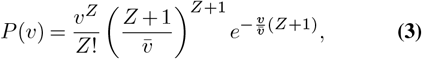

where 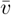 is the mean of the speed distribution and the vari-ance 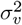 of *P* (*v*) is related to *Z* via 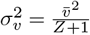. Each swim-ming speed data point in this work corresponds to the average of the DDM speeds measured in 2-5 movies, collected on one sample in a given condition, before the sample turned anaerobic. The anaerobic transition is readily identified in post-analysis by a sudden drop in swimming speed [37], and movies recorded past this transition were excluded. Error bars correspond to the standard deviation of the mean of these 2-5 measurements.

For the viscosity measurements, the diffusivity *D* of freely diffusing spheres (see Differential dynamic microscopy sub-section) was obtained by using the following intermediate scattering function in DDM analysis [84]:

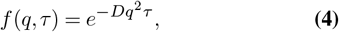

where *q* is the wave vector and *τ* the time. Viscosities were calculated using the Stokes-Einstein relation from the measured diffusivities.

#### Membrane potential quantification

To quantify PROPS fluo-rescence, only images recorded with 1 s exposure time were analysed. 50×50 pixels^2^ ROI were manually selected around cells that appeared flat in phase contrast images. The background intensity was subtracted for each cell by fitting a gaussian function to the intensity histogram of their respective ROI, and subtracting the mean of the fitted function. Cells were automatically segmented after background subtraction. First, pixels with intensities greater than one standard deviation of the pixel intensity distribution within the ROI were selected, and holes within the thresholded object were filled using the *imfill* function of MATLAB R2022a. This first thresholding step often included a ring of background pixels around the cells. A second threshold was applied to remove those pixels. Pixels with intensities greater than half the mean intensity of the 50% brightest pixels obtained after the first thresholding step were selected. These pixels defined the cell, and their average fluorescence was extracted.

For TMRM measurements, we followed the analysis method developed in [33], with minor adjustments. Cells were automatically segmented without background subtraction, first selecting all connected pixels with intensities one standard deviation above the mean intensity of the 50×50 pixels^2^ region around each cell. This step selected the cell body and a ring of background pixels around it. The cell body was segmented by only keeping pixels with intensities greater than the median value of all pixels selected after the first step. The intensities of the cells (*F*_in_) and background (*F*_out_) were measured by averaging, respectively, the intensities of all segmented pixels and of a 5×5 pixels^2^ area nearby each cell. We subtracted from *F*_in_ and *F*_out_ the intensity not caused by TMRM. For this, the average intensities of images recorded in the same tunnel slide filled with TMRM-free buffer were measured, before introducing cells and TMRM, at 9 different exposure times (20 ms - 200 ms). An affine function was fitted and used to estimate the value of the TMRM-independent background in experiments conducted at 50 ms exposure time.

The effect of the microscope’s point spread function (PSF) was accounted for and corrected as in [33], Fig. SI 18. Briefly, the 3-dimensional PSF was measured with 55 nm fluorescent beads (0.05um YG fluoresbrite, Polysciences)embedded in a 2% low melting point agarose gel. Using the PSF, simulated images of TMRM-loaded cells were produced as in [33], for different membrane voltage values and, thus, different cytoplasmic TMRM concentrations. Plotting the ratio of the simulated cell’s fluorescence (*F*_in_) to the background (*F*_out_) as a function of the known inner (*C*_in_) to outer (*C*_out_) TMRM concentration ratio returned a correction factor *S* given by 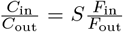, Fig.SI 18. The membrane voltage of cells measured in experiments was calculated as 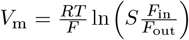. We did not correct for the fluorescence produced by membrane-bound TMRM. This voltage-independent fluorescence represents most of the signal in the LB/MM9 experiment, and, combined with the correction faclated in these conditions despite the unsuccessful TMRM loading.

### Data normalisation and visualisation

#### Motor speed normalisation and visualization

The results presented in Fig. 1A are calculated from the speed distribuions of Fig. SI 1, and those of Fig. 1B and C from Fig. SI 2. Motor speeds for each bead size were normalised by the average speed of all motors recorded with the corresponding bead size in the absence of glucose, Fig. 1A, and in the absence of butanol, Fig. 1B. The same was done for strain KAF95 in Fig. SI 14C using the speed distributions of Fig. SI 14D.

We used the absolute values of both CW and CCW speeds for all single-motor experiments, except in Fig. SI 14A and B, where we plot them separately using the normalisation method of Fig. 1A. The direction of the rotation does not affect the observed result.

In Fig. 2B, a dataset was generated for each treatment method, using the results of Fig. SI 3 (butanol), Fig. SI 4A (glucose, before normalisation, see paragraph about Fig. SI 4 below), and Fig. SI 4B(CCCP, before normalisation). Each butanol and glucose point is obtained from the mean absolute speeds of 0.5 and 1.5 µm beads on a given cell calculated for each 10-s (butanol) or 40-s (glucose) video. Each CCCP point is given by the mean absolute speeds of 0.5 and 1.5 µm beads on a given cell in a 3-s sliding window in Fig. SI 4B. For each dataset, the speeds of individual motors were normalised by the maximum motor speed recorded for the corresponding bead size in that particular dataset. For example, for the treatments that reduce PMF, this will be by the maximum speed recorded prior to the treatment. And for the glucose experiment, this will be the maximum speed recorded after addition of glucose.

In Fig. 2D, similar was done for butanol and glucose points using the results of Fig. SI 1 and Fig. SI 2, but here the average speeds were normalised by the maximum average speed of the data set, i.e. speed after glucose addition or before butanol treatment. Finally, for the light damage experiment, the single cell time traces of 2-4 cells per condition in Fig. SI 4C and D (before normalisation) are averaged and the mean absolute speed for 0.5 and 1.5 µm bead is calculated for a 10-s sliding window with 1 min step. The points in Fig. 2D are given by the 0.5 and 1.5 um bead speeds calculated in a given condition in a given 10-s window. The speeds of either bead recorded at various light powers were normalised by the maximum speeds recorded for the corresponding bead size across all cells before the light source was turned on.

In Fig. SI 4, motor speed time traces were normalised by the initial speed of each motor. Fig. 2 uses absolute speeds from the dataset of Fig. SI 4 before this normalisation.

In Fig. SI 5, for each type of treatment, the speed of each motor was normalised by the maximum speed recorded for that particular motor.

In Fig. SI 16G, the cell body and bead speeds of Fig. SI 16F are normalised by their initial values (average of first 5 points, tor, explains why a non-zero membrane voltage was calcu-^*t <*^ 8.2 s), before dissipation of PMF by CCCP.

In Fig. SI 19B, the motor speeds of each data set presented in Fig. SI 19A are normalised by the average speed of the respective data set at 0% butanol.

#### Motor torque normalisation and visualization

In Fig. 3, Fig. 4C, and Fig. SI 13, torques are calculated as described in the Data analysis section, using mean values of the speed distributions recorded for each bead size and butanol concentration in Fig. SI 2. Relative PMF values are calculated from the butanol concentrations and the calibration curve of Fig. SI 11.

In Fig. SI 17, indole concentrations used in [65] were converted to relative PMF values using the calibration curve from Krasnopeeva *et al*.: 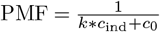, where *c*_ind_ is indole concentration in mM, and *k* = 2.58 mM^−1^ and *c*_0_ = − 0.365 are fitted constants [39]. Absolute torques were first calculated from the speeds and bead sizes taken from [65], and were normalised by the maximum torque obtained from this data set, *T*_max_=1630 pN nm.

#### Swimming speed normalisation and visualization

In Fig. 4, swimming speeds were normalised by the value measured at 0% butanol, with or without 8% Ficoll. The absolute swimming speeds measured in these experiments are presented in Fig. 4 *Inset*. In Fig. SI 12, speeds from the same data set are normalised by the values obtained at 0.25% butanol, with or without Ficoll.

#### Membrane potential normalisation and visualization

Normalised PROPS intensities are obtained, for each cell, by dividing the measured fluorescence with that measured on the same cell at 0% butanol, Fig. SI 7, or before glucose addition, Fig. SI 8.

## ACKNOWLEDGEMENTS

We would like to thank all members of the Pilizota, Poon, and Lo laboratories, especially Chao-Kai Tseng and Diana Coroiu, the latter for sharing EK01 Δ*cheY* strain. We thank Alexander Morozov, Calin Guet, and Jochen Arlt for their support and useful discussions. TP, EK, and CJL were supported by the Human Frontier Science Program grant (RGP0041/2015), EK was supported by the EMBO Long-Term Fellowship ALTF 44-2021 and the Scottish Universities Physics Alliance (SUPA) Short-Term Visit Funding Scheme to CJL’s lab. TP, LLN and UEBP were supported by the Leverhulme Trust grant RPG-2019-187 and the EPSRC established career fellowship EP/V03264X/1. LLN was supported by a Research Fellowship from the Royal Commission for the Exhibition of 1851. CJL was supported by NSTC Taiwan (NSTC-109-2628-M-008-01-MY4), and UEBP was supported by a CONAHCyT Mexico PhD scholarship.

## AUTHOR CONTRIBUTIONS

EK, LLN, and TP designed the experiments. EK, LLN, and UEBP performed the experiments, and EK and LLN carried out the analysis with input from CJL, WP, and TP. The interpretation of the results was performed by EK, LLN, CJL, WP, and TP. EK, LLN, and TP cowrote the paper and all authors commented on the results and the manuscript.

## Supplementary Information

### Supplementary Tables. Tables SI 1 - SI 8

**Table SI 1.** Summary of data set sizes for single-cell, single-motor experiments.

| Experiment | Figures | Number of cells |  |  |  |  |
| --- | --- | --- | --- | --- | --- | --- |
| | | 0.35 $\mu\text{m}$ | 0.5 $\mu\text{m}$ | 0.75 $\mu\text{m}$ | 1 $\mu\text{m}$ | 1.5 $\mu\text{m}$ |
| No glucose | 1A, 2D, SI 1, SI 14A-C | 25 | 27 | 24 | 27 | 22 |
| 0.3% glucose |  | 37 | 30 | 27 | 24 | 22 |
| Butanol 0% | 1B-C, 2D, 3, 4A-C, SI 2, SI 11, SI 13, SI 17, SI 19, SI 21 | 78 | 32 | 80 | 73 | 57 |
| Butanol 0.1% |  | 51 | 21 | 36 | 69 | 16 |
| Butanol 0.25% |  | 50 | 11 | 42 | 51 | 77 |
| Butanol 0.5% |  | 29 | 16 | 49 | 115 | 30 |
| Butanol 0.75% |  | 33 | 12 | 40 | 119 | 39 |
| Butanol 1% |  | 32 | 6 | 29 | 104 | 27 |
| Light damage 333 mW/cm<sup>2</sup> | 2D, SI 4C-D |  | 3 |  |  | 4 |
| Light damage 2338 mW/cm<sup>2</sup> |  |  | 2 |  |  | 3 |
| KAF95 No glucose | SI 10C, SI 14C-D |  | 28 |  |  | 34 |
| KAF95 0.3% glucose |  |  | 31 |  |  | 33 |

**Table SI 2.**
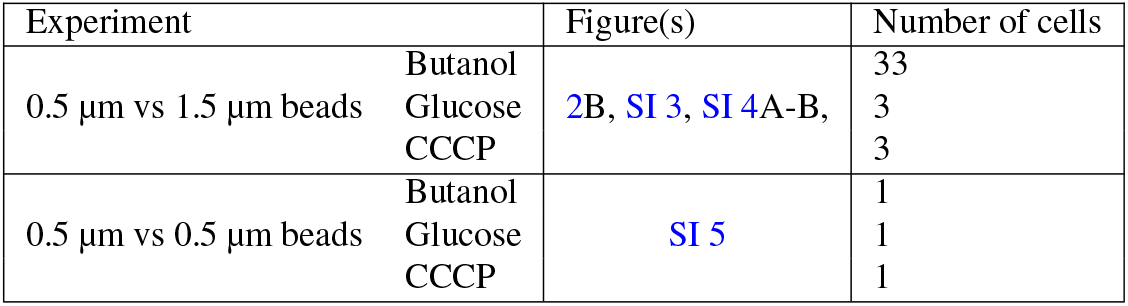
Summary of data set sizes for single-cell, double-motor experiments.

**Table SI 3.** Summary of data set sizes for membrane potential measurements with PROPS.

| Experiment | Figure(s) | Number of cells |
| --- | --- | --- |
| TMRM, BMB | SI 6 | 79 |
| TMRM, MM9 | SI 6 | 60 |
| PROPS, butanol and CCCP treatment | SI 7 | 156 |
| PROPS, glucose addition | SI 8 | 42 |

**Table SI 4.** Time scales of PMF change experiments across different works compared to stator dynamics. PMF in this work has been changed with various methods within 2-30 min, covering the range of previous work of Gabel and Berg [29]. Although butanol causes PMF changes within 2 sec [39], in this work we recorded only the steady-state motor speeds, 10-15 minutes after the flush. In Fung and Berg’s pioneer work demonstrating PMF-speed proportionality with external voltage source, voltage changed rapidly and stayed at a same value for only about 4-8 s. A single stator unit has been estimated to take 10-80 s for association with a motor. Thus, Fung and Berg experiments were most likely probing motors with fixed number of stator units (PMF change time < stator unit association time), whereas in Gabel and Berg’s work and in this work the number of stator units is likely varied, allowing the motor to adapt its new steady state at a new PMF level (PMF change time > > stator unit association time).

| Experiment | Time scales | References |
| --- | --- | --- |
| Glucose | 30-40 min | This work, Fig. SI 4A |
| Butanol | 10-15 min | This work |
| CCCP | 2-2.5 min | This work, Fig. SI 4B |
| Light | 3.5-10 min | This work, Fig. SI 4C-D |
| CCCP/sodium azide | 10 min | [29, 58] |
| External voltage | 4-8 sec | [24, 57] |
| **Stator unit association time** | 20 s | [24] |
|  | 20 s | [13] |
|  | 20-80 s | [69] |
|  | 10-40 s | [85] |

**Table SI 5.** Data set sizes obtained in the geometry of ref. [29].

| Experiment | Figure | Medium | Bead size | Number of cells |
| --- | --- | --- | --- | --- |
| Passive vs. active rotation statistical analysis | SI 15 | MB | 1.5 | 18 |
|  | SI 15 | MB | 0.75 | 13 |
|  | SI 15 | MBG | 0.5 | 34 |
|  | SI 15 | GMBG | 0.5 | 85 |
| CCCP, passively rotating cell | SI 16A-E | MBG, GMBG | 0.5 | 4 |
| CCCP, actively rotating cell | SI 16F-G | MBG | 0.5 | 1 |

**Table SI 6.** Summary of bacterial strains used in this study.

| Strain | Genotype / Description | Relevant Figures | Reference |
| --- | --- | --- | --- |
| EK01 | MG1655 derivative with *fliC* replaced by *fliC<sup>sticky</sup>* | Fig. 1, Fig. 2, Fig. 3, Fig. SI 1, Fig. SI 2, Fig. SI 3, Fig. SI 4 A&B, Fig. SI 5, Fig. SI 11, Fig. SI 14 | [39] |
| EK01 $\Delta tolC$ | EK01 derivative with *tolC* deletion | Fig. SI 6 | [41] |
| EK07 | EK01 derivative with chromosomally integrated pHluorin | Fig. 2, Fig. SI 4 C&D | [39] |
| KAF95 | RP437 $\Delta fliC$ $\Delta cheY$ with plasmid encoding *fliC<sup>sticky</sup>* | Fig. SI 14 | [29] |
| YD133+pWR20 +pBAD/PROPS | Keio collection derivative with $\Delta fliC$, $\Delta pilA$, $\Delta flgE$ with pWR20 plasmid constitutively expressing cytoplasmic GFP (KanR) and pBAD expressing arabinose-inducible PROPS (AmpR) | Fig. SI 7, Fig. SI 8, Fig. SI 9, Fig. SI 10 | [77, 78]<br>[43] |
| AD80 | Smooth swimming MG1655 $\Delta cheY$ | Fig. 4, Fig. SI 12 | [37] |
| EK01 $\Delta cheY$ | EK01 derivative with motors rotating exclusively counterclockwise | Fig. SI 15, Fig. SI 16, Fig. SI 24, Fig. SI 25, Videos SI 3 and SI 4 | From Diana Coroiu |

**Table SI 7.** Media used in this work. pH was adjusted to 7.0-7.2 in all experiments except for light damage experiments, where pH=7.5, and pH values are also given throughout *Materials and Methods*.

| Medium | Acronym | Composition |
| --- | --- | --- |
| Lysogeny broth | LB | 10 g tryptone, 5 g yeast extract, 10 g NaCl per 1 L |
| Tryptone broth | TB | 10 g tryptone, 5 g NaCl per 1 L |
| Modified M9 | MM9 | 50 mM Na<sub>2</sub>HPO<sub>4</sub>, 25 mM NaH<sub>2</sub>PO<sub>4</sub>, 8.5 mM NaCl and 18.7 mM NH<sub>4</sub>Cl with added 0.1 mM CaCl<sub>2</sub>, 1 mM KCl, 2 mM MgSO<sub>4</sub> |
| Motility buffer | MB<br>MBG | 10 mM potassium phosphate, 67 mM NaCl, 0.1 mM EDTA<br>MB+0.3% glucose |
| Gabel's motility buffer | GMB<br>GMBG | 10 mM potassium phosphate, 0.1 mM EDTA, 1 mM L-methionine<br>GMB+0.3% glucose |

**Table SI 8.**
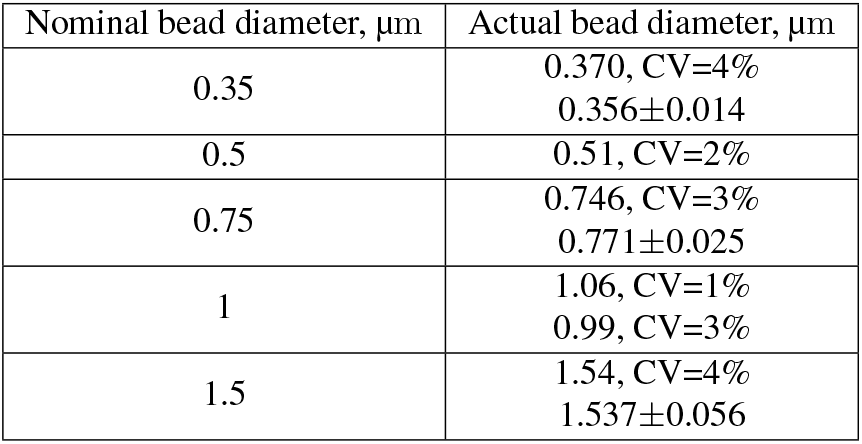
Bead diameters used in the experiments.

## Supplementary Figures

Figures re given in the order mentioned in the main text, followed by mentions in the Supplementary Information (SI) text.

**Figure SI 1.**
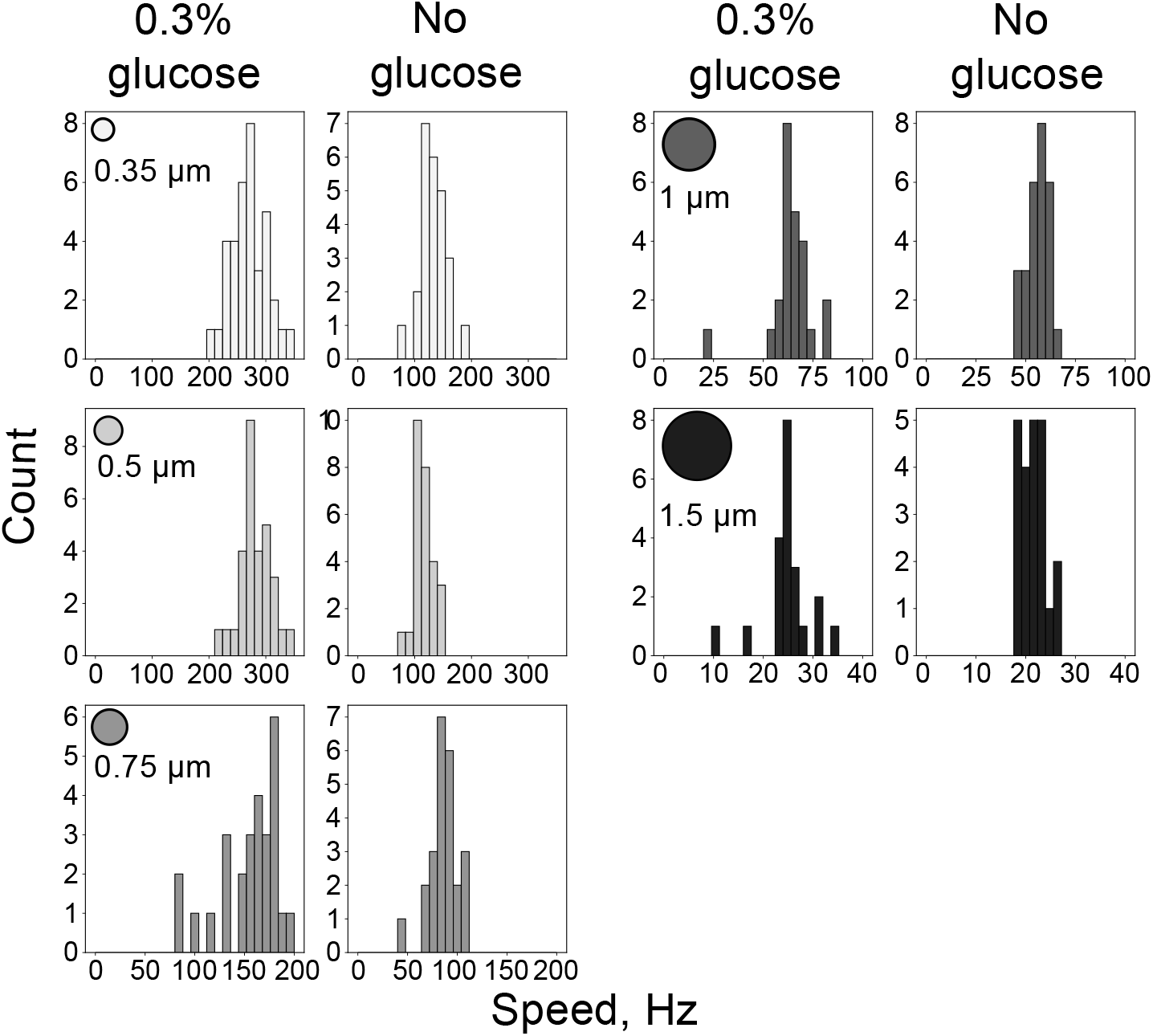
Motor speed distributions, supplementary to Fig. 1A. Histograms of motor speeds for each experiment with and without glucose (25 bins are used for a given range). Bead colour coding is the same as in the main text. See Table SI 1 for sample sizes.

**Figure SI 2.**
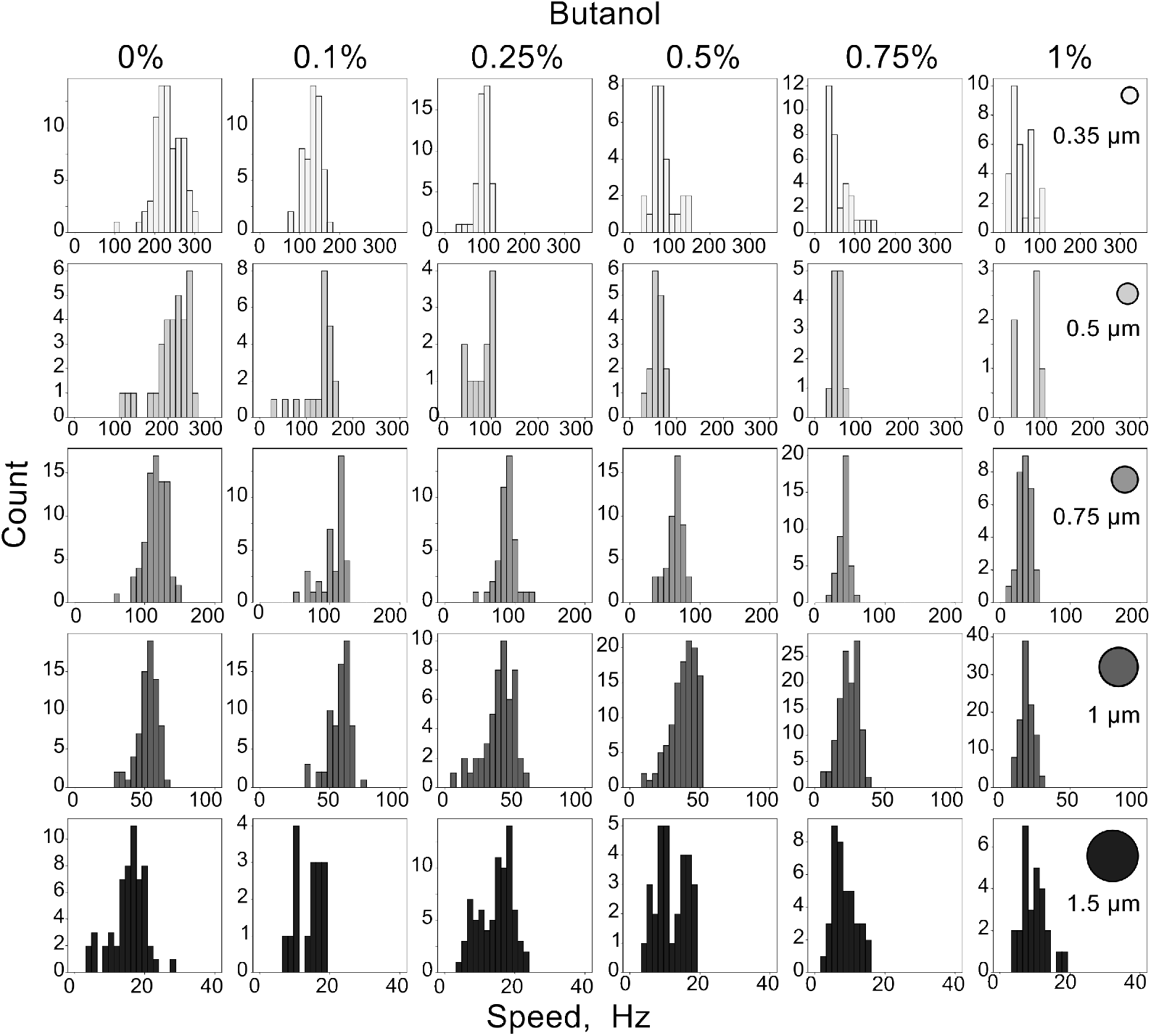
Motor speed distributions, supplementary to Fig. 1B and C. Histograms of motor speeds for all butanol experiments (25 bins are used for a given range). Bead colour coding is the same as in the main text. See Table SI 1 for sample sizes.

**Figure SI 3.**
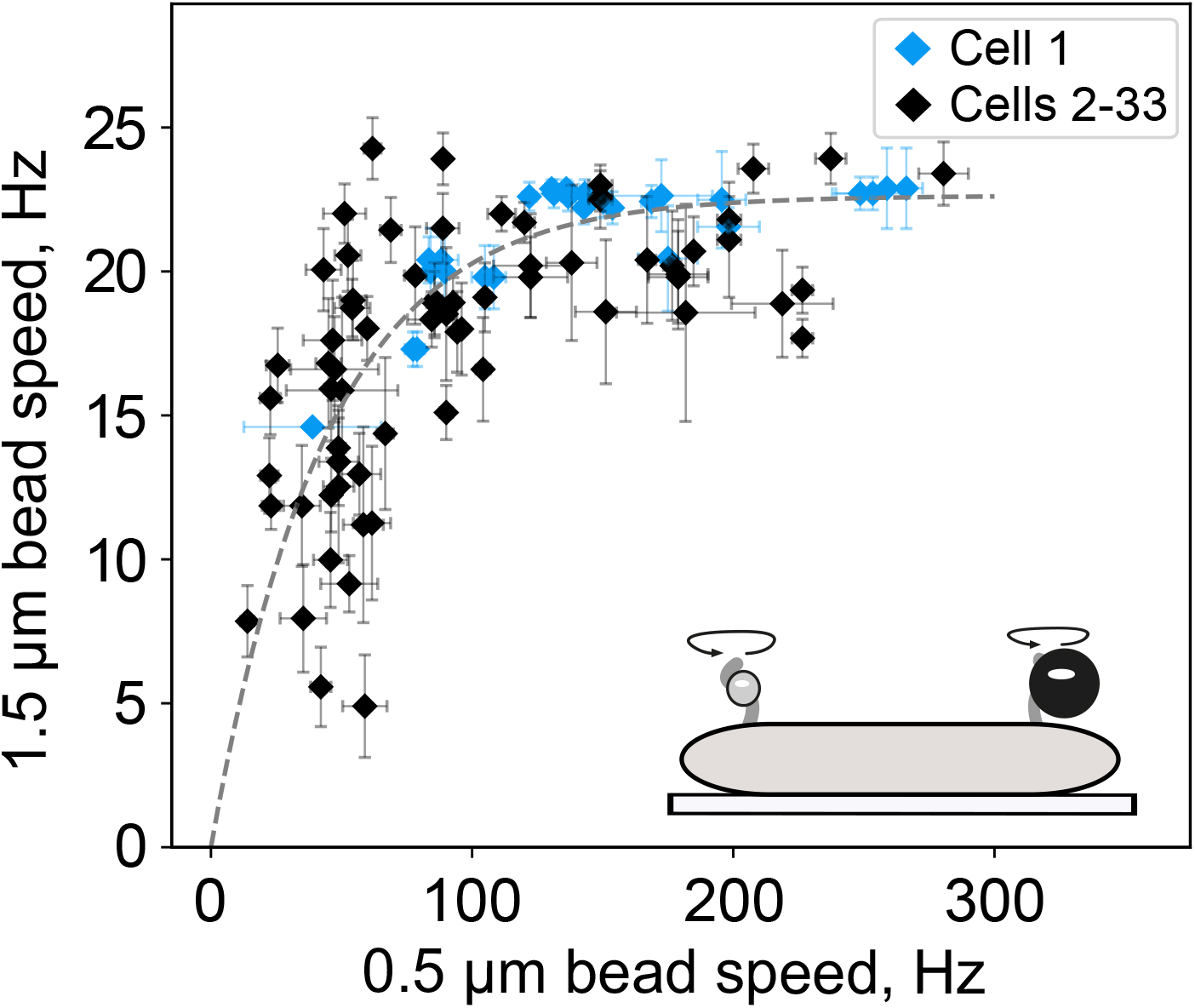
Absolute speed of two motors belonging to one cell under different loads is not linearly proportional at different butanol concentrations, supplementary to Fig. 2. Filamentous cells treated with various butanol concentrations. Most of the time, one cell was treated with one or two butanol concentrations (black markers), whereas the blue points show a cell that was treated with four butanol concentrations with multiple movies recorded for each concentration. Dashed line: exponential fit applied to all the data, 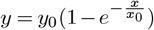, where *y*_0_ = 21.246 Hz and *x*_0_ = 38.726 Hz are fitted parameters. The error bars show the standard deviation of the single bead speeds, and Table SI 2 sample sizes.

**Figure SI 4.**
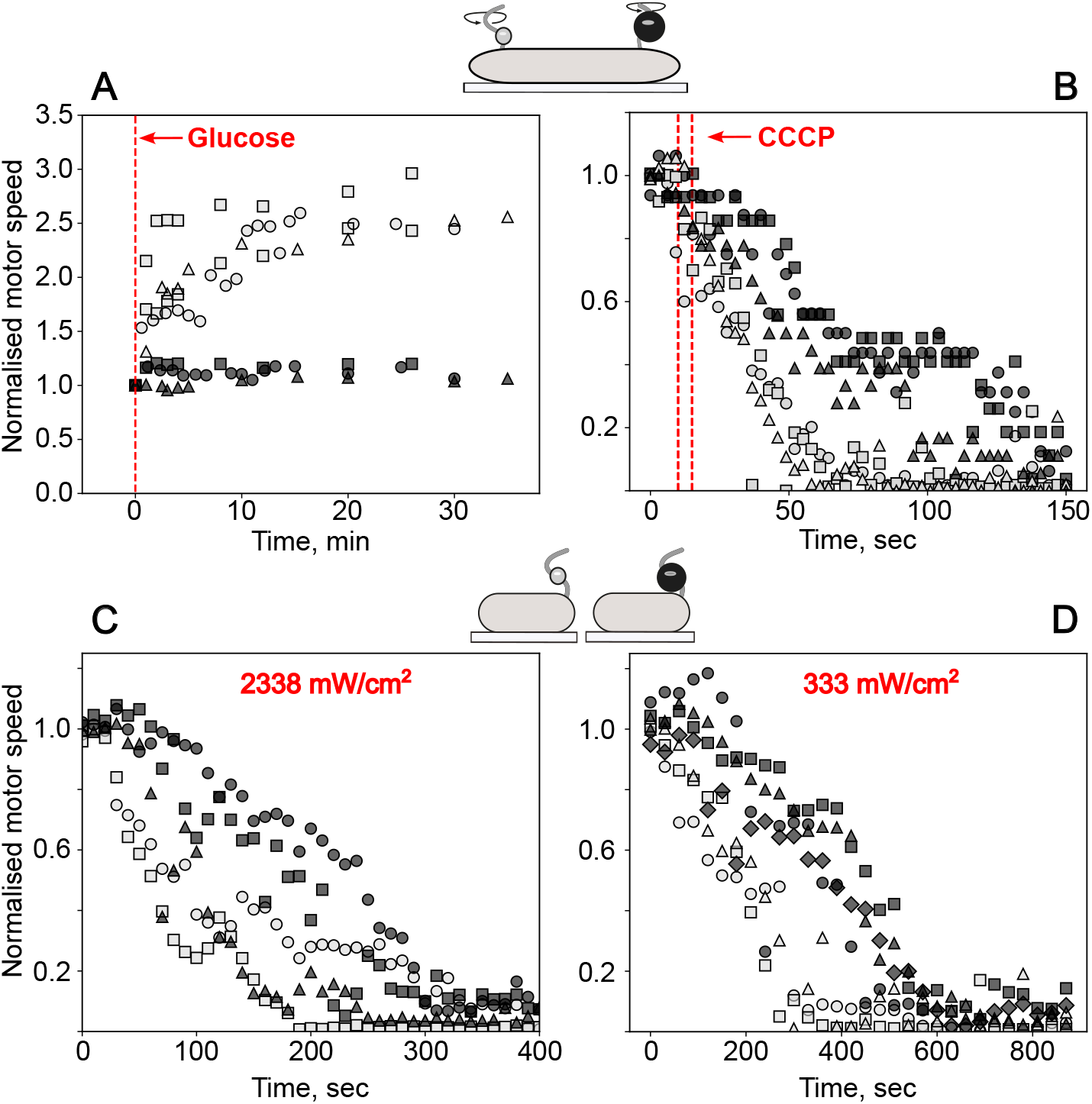
Dynamics of the speed changes for motors under different loads, supplementary to Fig. 2. As in the main text, dark grey markers show 1.5 µm beads, and light grey 0.5 µm. Different symbol types depict different independent experiments. **A**. and **B**. Time-speed traces of the double-bead experiments where filamentous cells are treated with glucose or CCCP, respectively (*Materials and Methods*). In **A**, cells are initially in MM9, and at 0 min the medium is replaced with MM9+0.3% glucose. In **B**, cells are initially in MM9+0.3% glucose, and CCCP is added at 10-15 s. Addition of CCCP or glucose is indicated with the dashed lines. In **B**, two such lines indicate the interval of addition of CCCP. **C**. and **D**. Time-speed traces of the single-motor experiments with cells exposed to 395 nm light of 2338 mW/cm^2^ or 333 mW/cm^2^ power. See *Materials and Methods* for acquisition and normalisation procedures and the use of these results in Fig. 2D, and Tables SI 1 - SI 2 for sample sizes.

**Figure SI 5.**
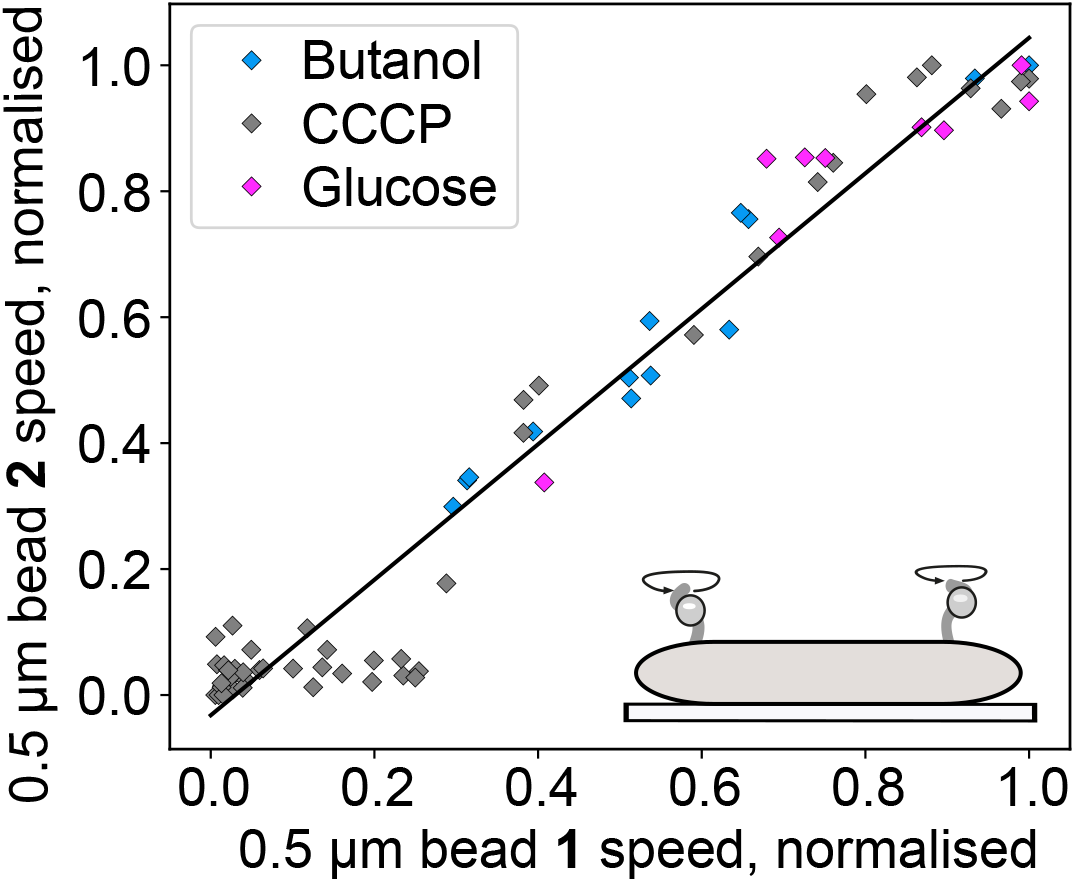
Speeds of two 0.5 µm beads on the same cell are linearly proportional. Double-bead experiments with filamentous cells having two beads of the same size (0.5 µm) attached. The cells are treated with butanol, CCCP, or glucose the same way as in Fig. 2 and Fig. SI 4. Each data set shows one cell with two 0.5 µm beads. Solid line: linear fit *y* = 1.0295*x*. See *Materials and Methods* for normalisation procedures.

**Figure SI 6.**
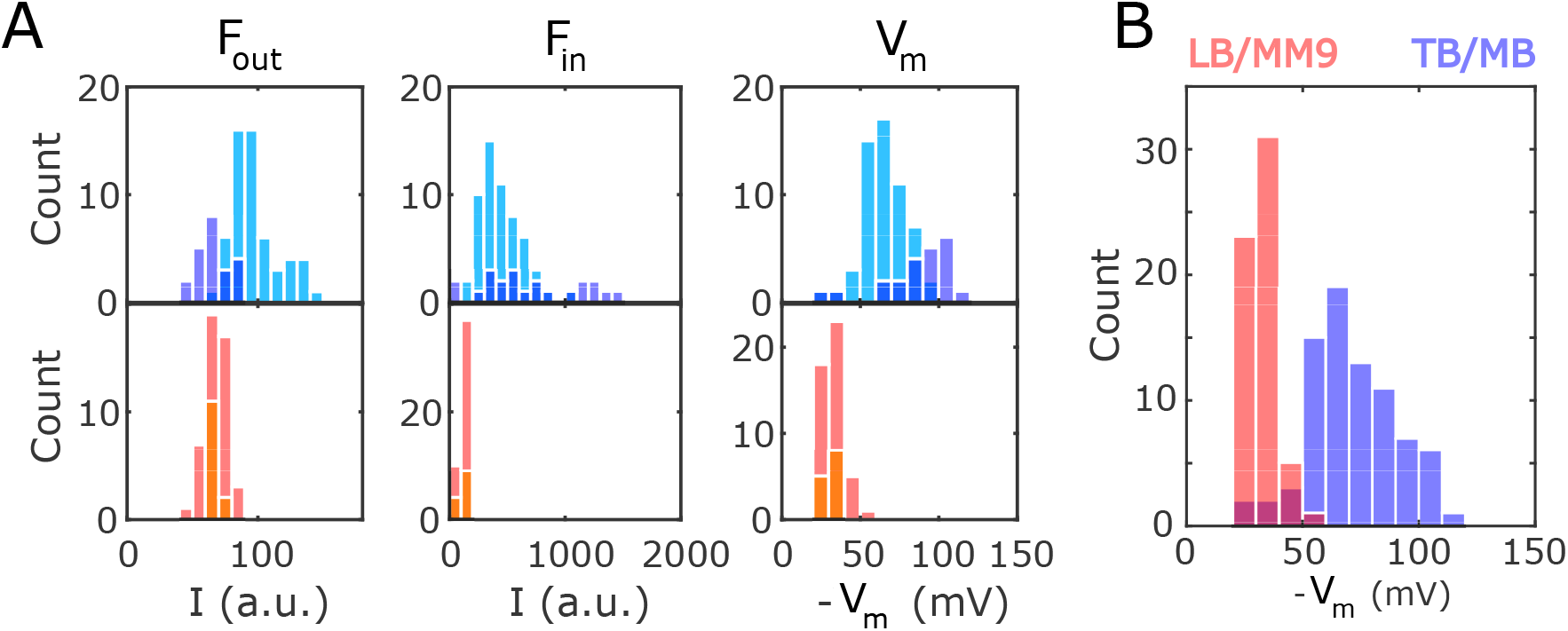
TMRM does not load in cells grown in LB and imaged in MM9. Cells grown in TB (blue) and LB (red) were incubated and imaged in MB and MM9, respectively, both supplemented with 100 nM TMRM. The TB/MB preparation replicates the conditions used in [33], whereas LB/MM9 corresponds to the main experimental condition used throughout this work. See *Materials and Methods* for details. **A**. Histograms of external TMRM intensity (*F*_out_), cytoplasmic TMRM intensity (*F*_in_), and calculated membrane voltage (*V*_m_) for each cell measured in each condition (*Materials and Methods*, Data analysis section). Light blue and red are cells imaged immediately (<4 min) after being introduced into the channel, and dark blue and red those imaged after 10 min (and after washing out unattached cells with fresh MB or MM9 both with 100 nM TMRM). LB-grown cells in MM9 are only marginally brighter than the background, indicating no loading (average *I*_cell_ - *I*_background_ = 47 for LB-grown cells in MM9, vs average *I*_cell_ - *I*_background_ = 430 for TB-grown cells in MB). **B**. Histogram of Vm calculated for all cells included in panel A. The non-zero but weak signal, and corresponding non-zero *V*_m_ value calculated for cells grown in LB and imaged in MM9, is attributed to TMRM staining the cells’ envelope, not to *V*_m_-dependent cell-loading [33]. See Table SI 3 for sample sizes.

**Figure SI 7.**
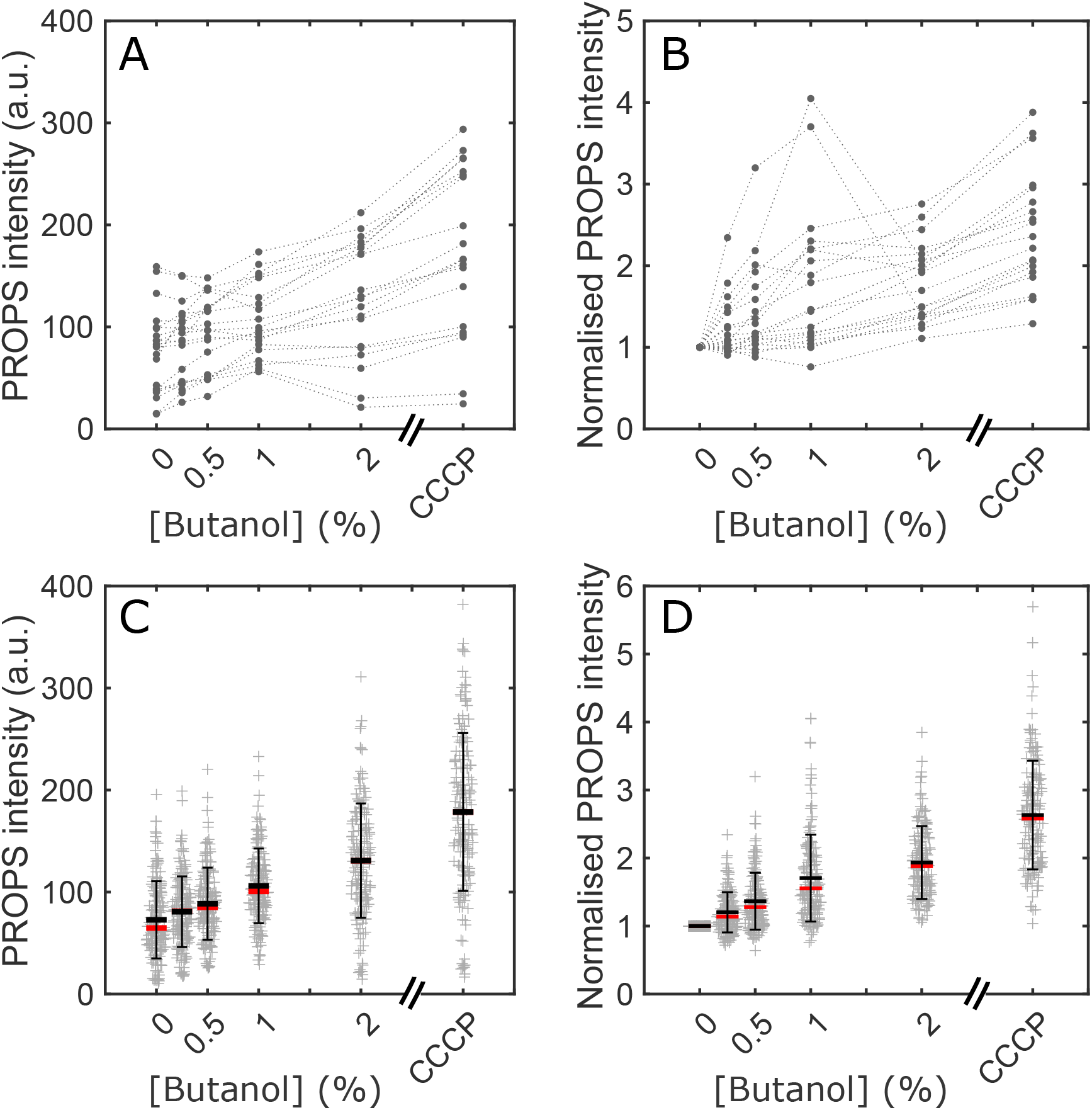
Measurements of relative *V*_m_ during CCCP and butanol treatment show expected decrease. Cells were first exposed to increasing butanol concentrations (0% - 2%), and then to ∼ 250 µM CCCP, both in MM9+0.3% glucose and in the same channel. Measurements were conducted in quick succession, with each butanol concentration and the CCCP measurement taking ∼ 10 min. **A. & B**.: absolute and normalised PROPS intensities for 20 randomly selected “flat” cells (see *Materials and Methods*). Dashed lines connect the points of each individual cell. **C. & D**.: absolute and normalised PROPS intensities for all measured cells (see Table SI 3 for all sample sizes). Black line shows the average, red the median, and the error bars are standard deviations. See *Materials and Methods* for normalisation procedures.

**Figure SI 8.**
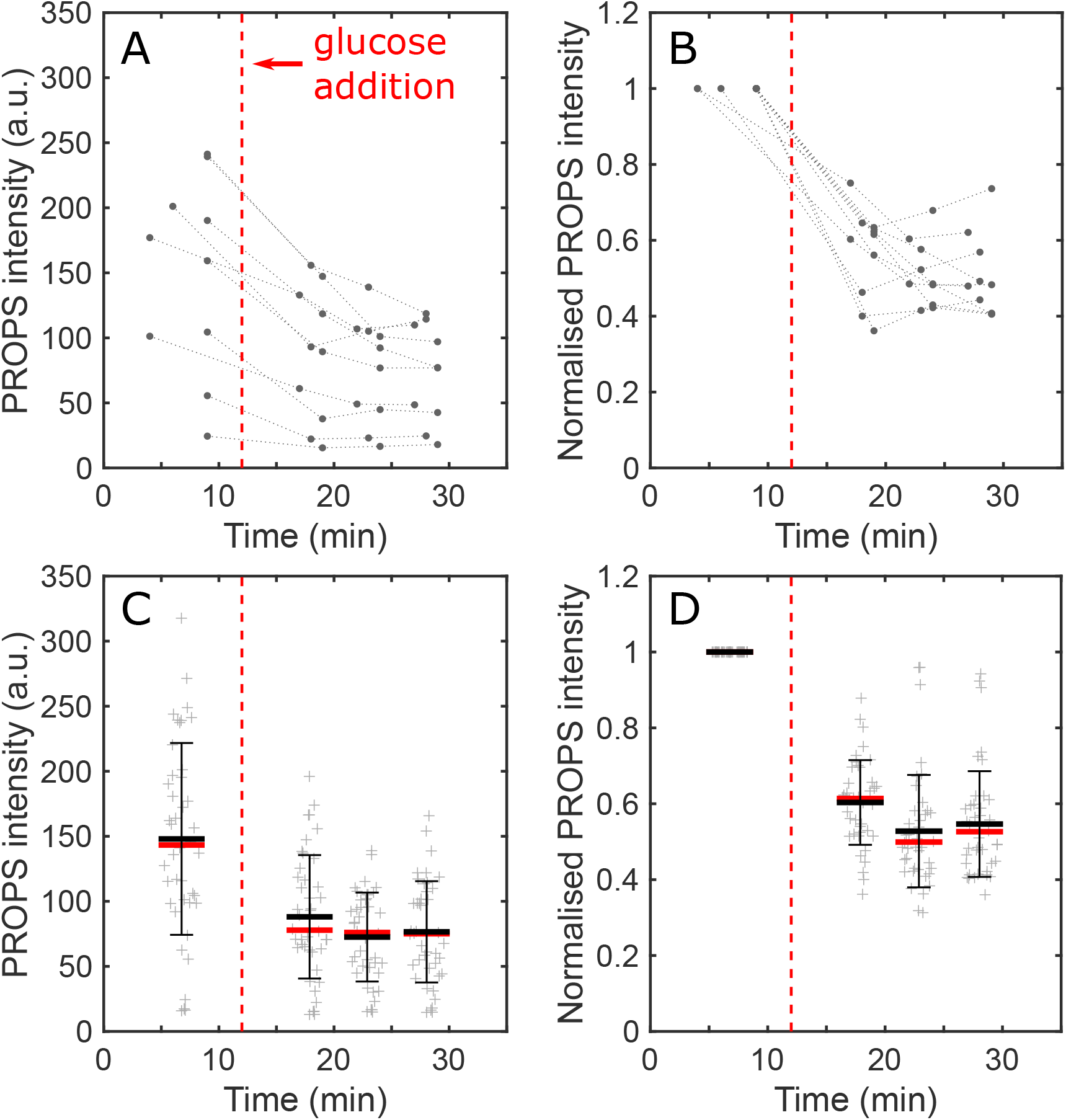
Measurements of relative *V*_m_ during addition of glucose show expected increase. First measurement was performed in MM9 (no glucose), approximately 10 min after cells were introduced into the tunnel slide. At *t* = 12 min (vertical red dashed line), MM9+0.3% glucose was flushed into the chamber. **A. & B**.: absolute and normalised PROPS intensities for 10 randomly selected ‘flat’ cells (see *Materials and Methods*). Dashed lines connect the points of each individual cell. **C. & D**.: absolute and normalised PROPS intensities for all measured cells (see Table SI 3 for all sample sizes). Black line shows the average, red the median and error bars are standard deviations. See *Materials and Methods* for normalisation procedures.

**Figure SI 9.**
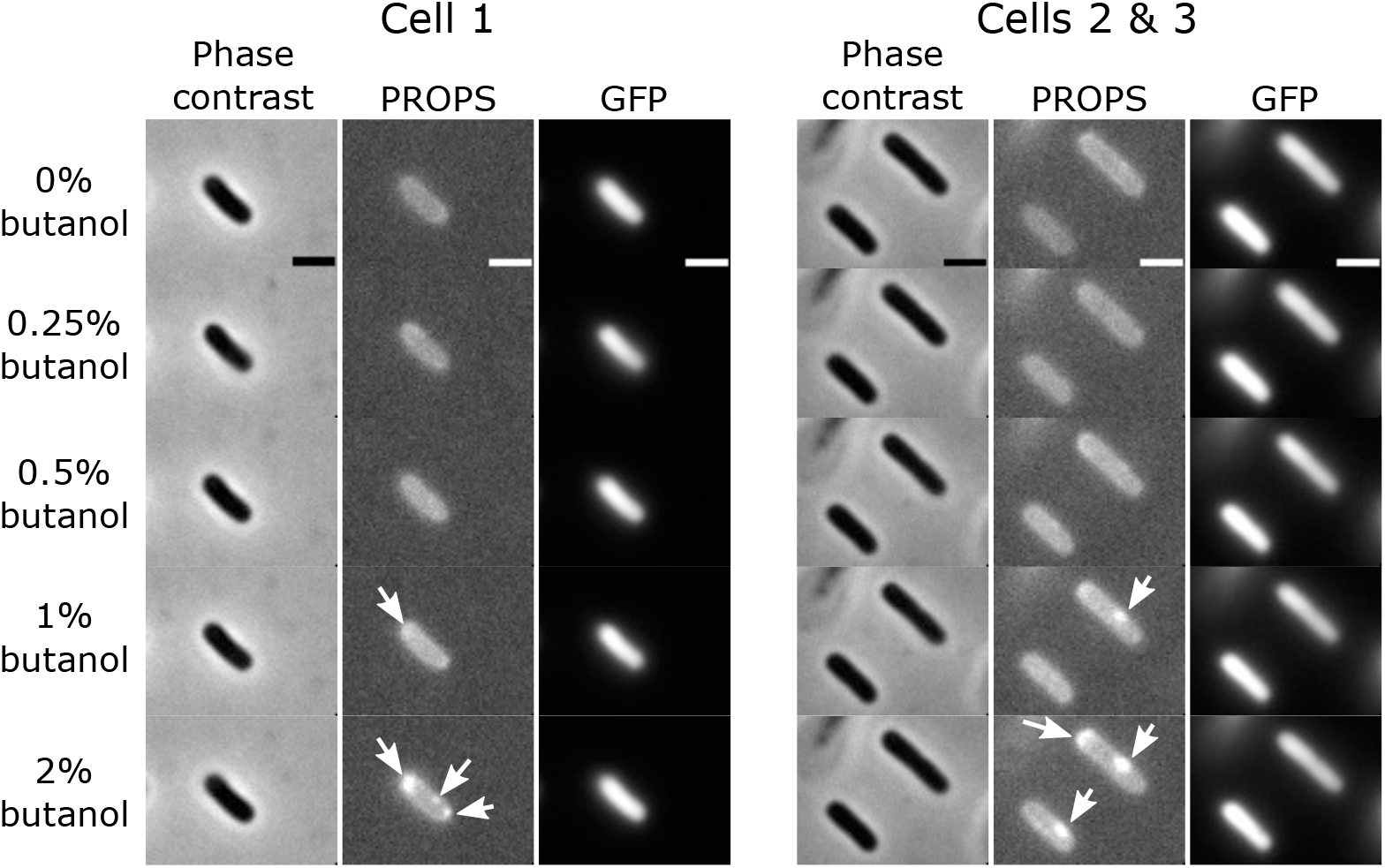
PROPS aggregates in cells at high butanol concentrations. Representative phase contrast, PROPS and GFP images recorded at different butanol concentrations in MM9+0.3% glucose. At 1% and 2% butanol concentrations, bright clusters of PROPS become visible (white arrows). GFP is not affected by butanol and remains homogeneously distributed in the cells. Scale bars: 2 µm.

**Figure SI 10.**
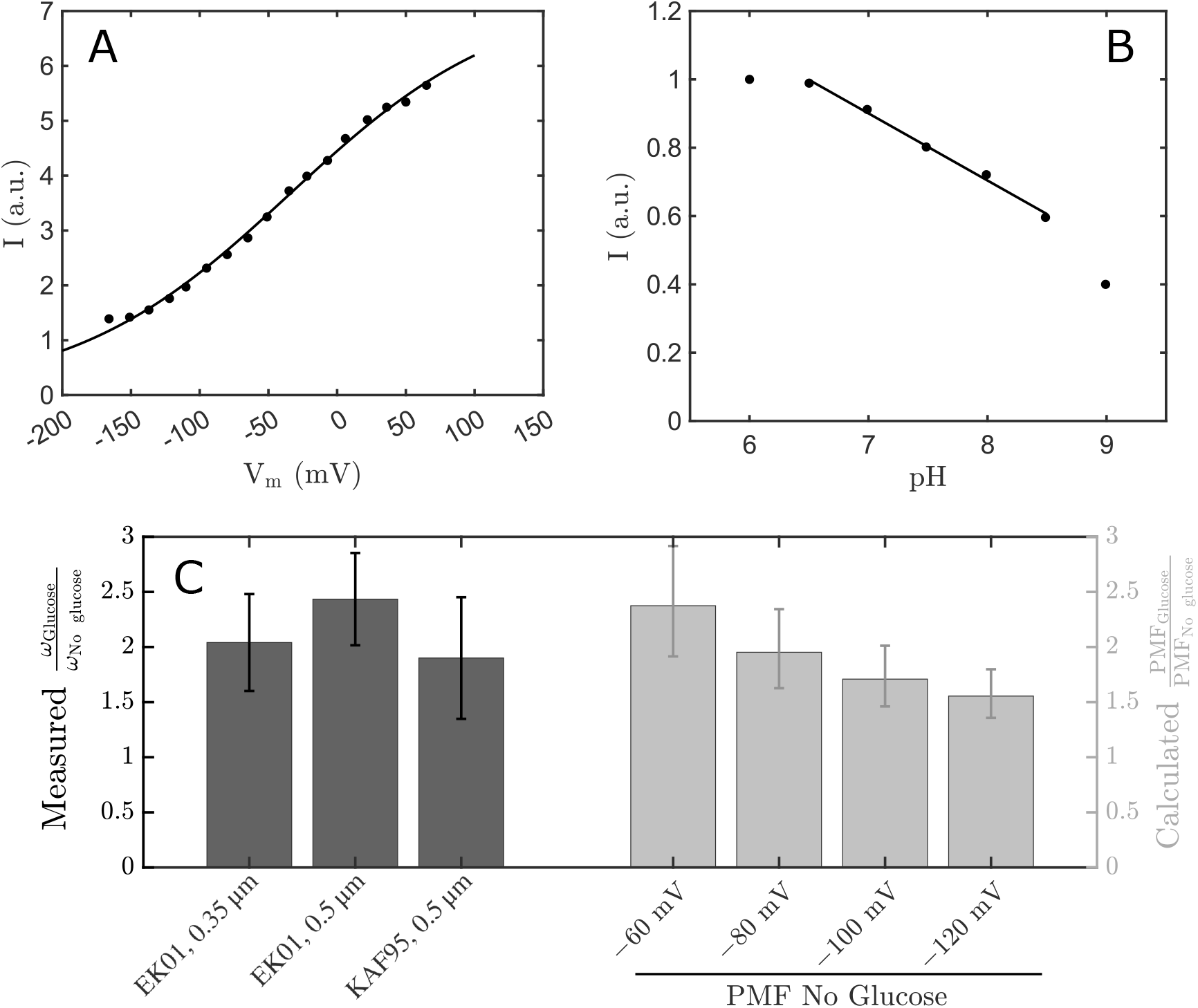
Calibration of *V*_m_ sensor PROPS, and comparison of relative motor speed increase with relative PMF change upon glucose addition. **A**. PROPS intensity as a function of externally applied membrane voltage at constant internal pH reproduced from [43], with a sigmoidal fit of the data obtained using a logistic model: 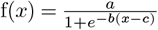. **B**. PROPS intensity as a function of internal pH at near-zero *V*_m_ (cells were treated with CCCP) reproduced from Supplementary Information of [43]. The line is an affine fit in the region of the graph relevant to the present study. **C**. Comparison of the glucose/no glucose low-load motor speed ratios (left vertical axis, black bars) recorded in EK01 and KAF95 strains with the glucose/no glucose PMF ratios calculated from PROPS measurements. 4 different assumptions for the value of PMF with no glucose are given (see SI text). Error bars for the glucose/no glucose low-load motor speed ratios are standard deviations. Error bars for the calculated PMF ratios are obtained from the standard deviation of normalised PROPS intensity in Fig. SI 8D, neglecting other uncertainties (see SI text).

**Figure SI 11.**
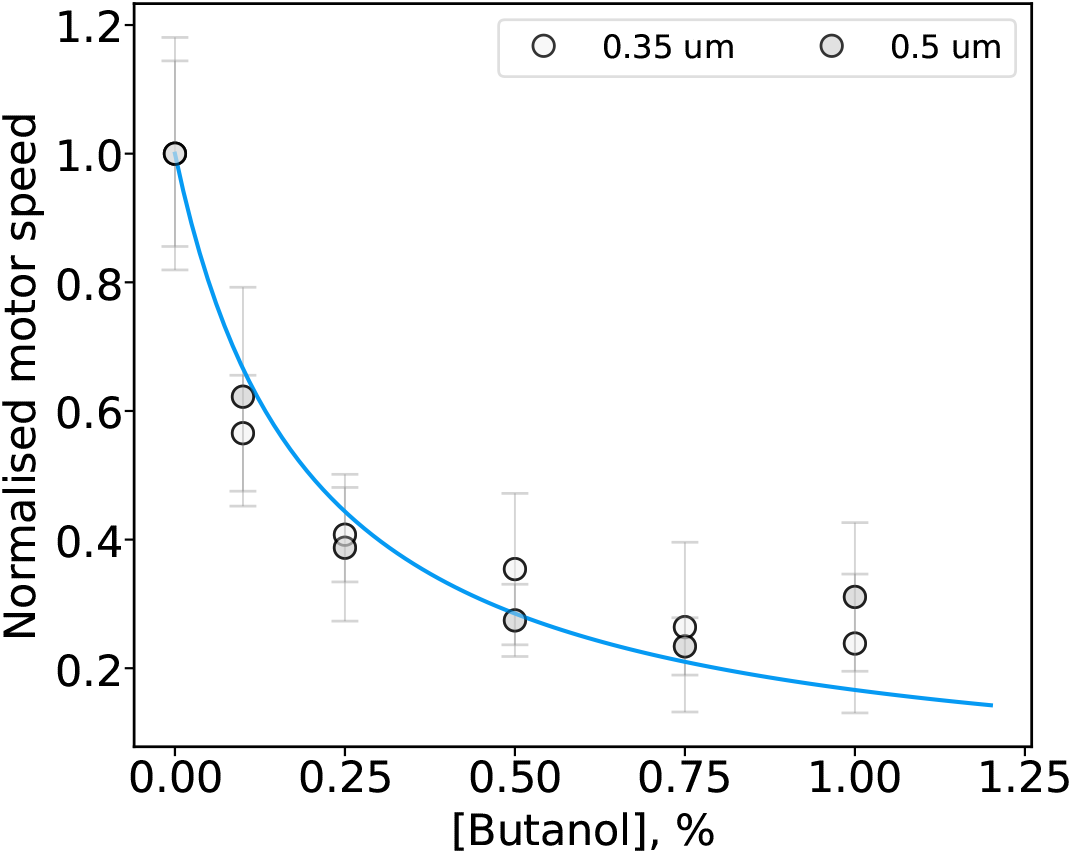
Calibration curve for converting butanol concentration to relative change in the PMF. Speed-butanol curve is fitted with a hyperbola 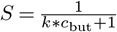, where is butanol concentration in volume/volume percentage, *S* is a relative motor speed, and *k* = 5.01 is a fitted constant. Since normalised motor speed of the BFM under low viscous load is assumed to be proportional to PMF, the equation for normalised PMF is also.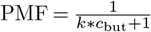 Data points are from Fig. 1B *Inset*. Error bars are standard deviations.

**Figure SI 12.**
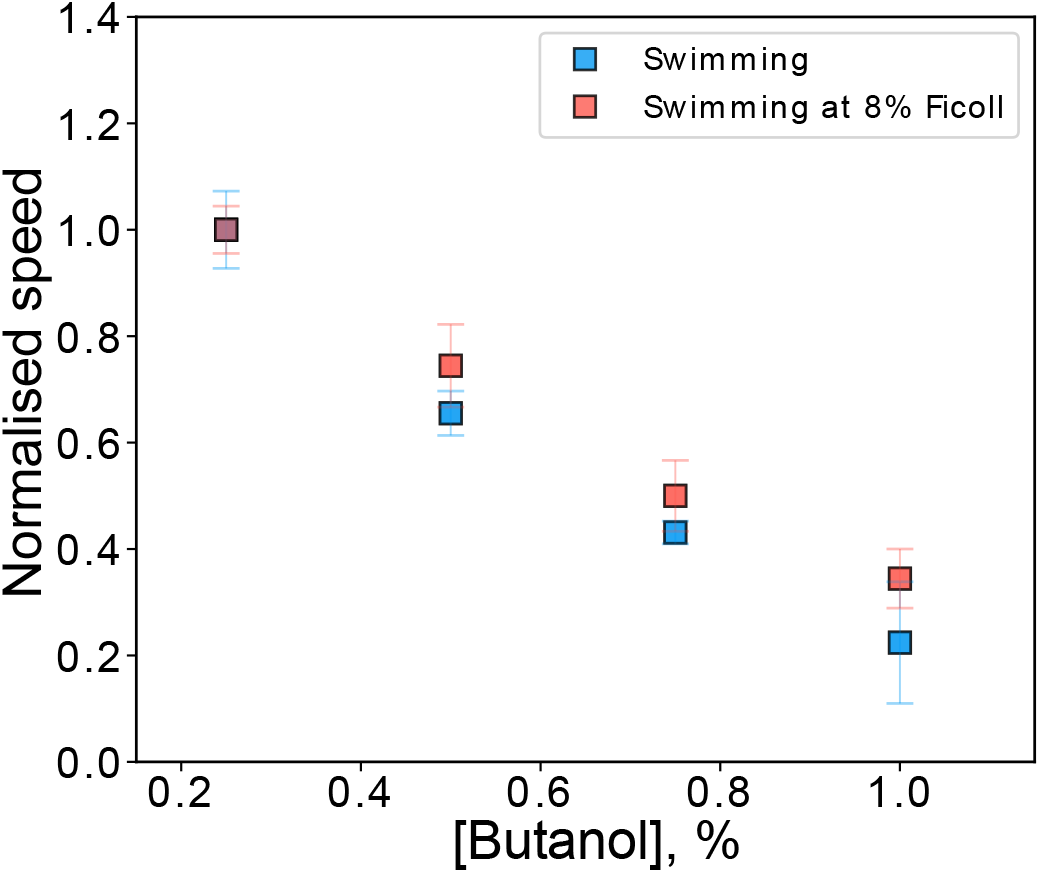
Free swimming speeds with and without Ficoll from Fig. 4B collapse to the same curve for low PMF values. Here, speeds are normalised to the speed at 0.25% butanol, corresponding to 44% of maximum PMF. Error bars are standard deviations. See *Materials and Methods* for normalisation procedures.

**Figure SI 13.**
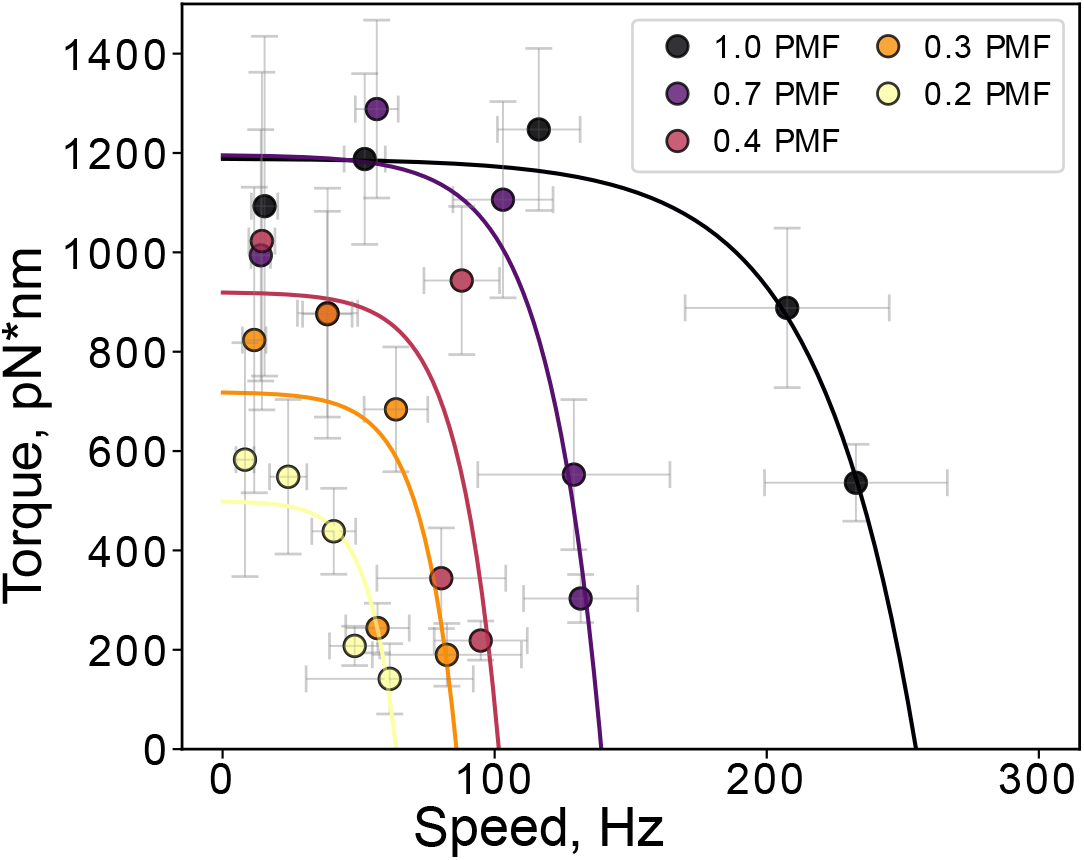
Torque-speed curves for data in Fig. 1C. Motor torque is calculated as described in *Materials and Methods*, and, as before, the PMF is scaled to PMF of a bacterium in MM9+0.3% glucose medium with no butanol added. The fits are exponential decay curves 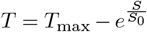, where *S* is a motor speed in Hz, and *S*_0_ is a free fitting parameter. Data points show mean with the standard deviation. See Table SI 1 for sample sizes.

**Figure SI 14.**
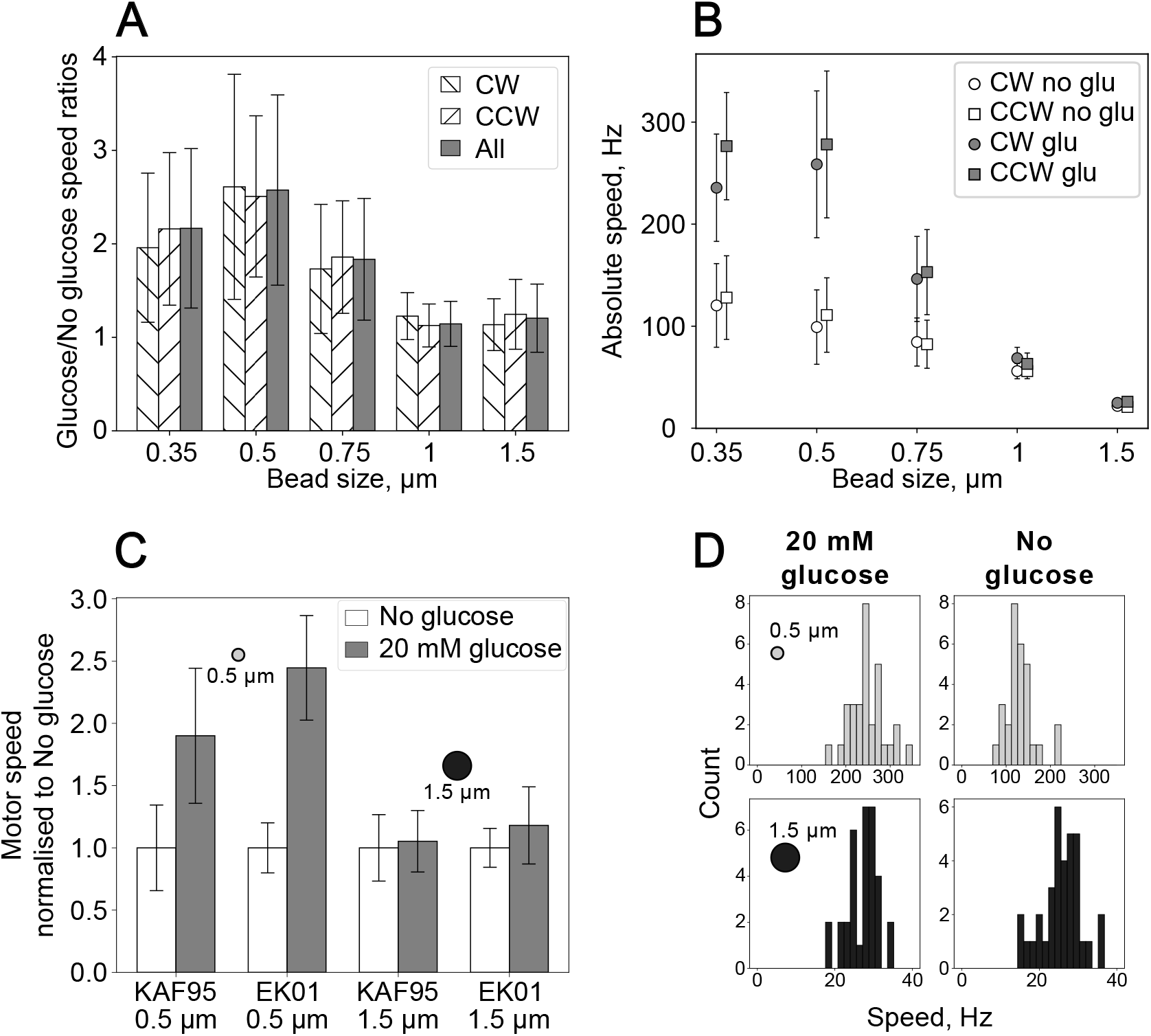
The torque saturation effect is independent of the direction of motor rotation and is reproducible with CheY deficient KAF95 strain (RP437 derivative). **A**. Data from the main Fig. 1A plotted separately for CCW (positive motor frequencies), or CW (negative motor frequencies) rotation, and for all absolute speeds (same as in Fig. 1A). The ratios between Glucose and No-glucose speeds are conserved irrespective of the rotational direction. **B**. Absolute speeds of BFM at different loads with glucose (grey) or without (white) are shown separately for CW (circles) and CCW (squares) rotation. As expected [3, 21, 86], CW speeds are lower than CCW in a fast rotation regime (≥ 100 Hz), and equal in slow. **C**. In this work we used EK01 strain, which is an MG1655 derivative with the wild type BFM, whereas previous work used the RP437 derivative, KAF95 lacking CheY and thus, with BFM capable of only CCW rotation [25, 29, 53, 54]. When grown in the same conditions, these strains have been reported to swim at somewhat different speeds [37]. Consistent with our findings, relative speed change for cells with and without glucose is different under 0.5 or 1.5 µm beads load but similar to that of the EK01 strain (MG1655 derivative) used in this work. **D**. Speed histograms for KAF95 speed data shown in C. Histograms are plotted in ranges (0,350) for 0.5 µm and (0,40) for 1.5 µm beads (25 bins are used for each range). Bead colour coding is the same as in the main text. Error bars are standard deviations in all panels. See Table SI 1 for sample sizes.

**Figure SI 15.**
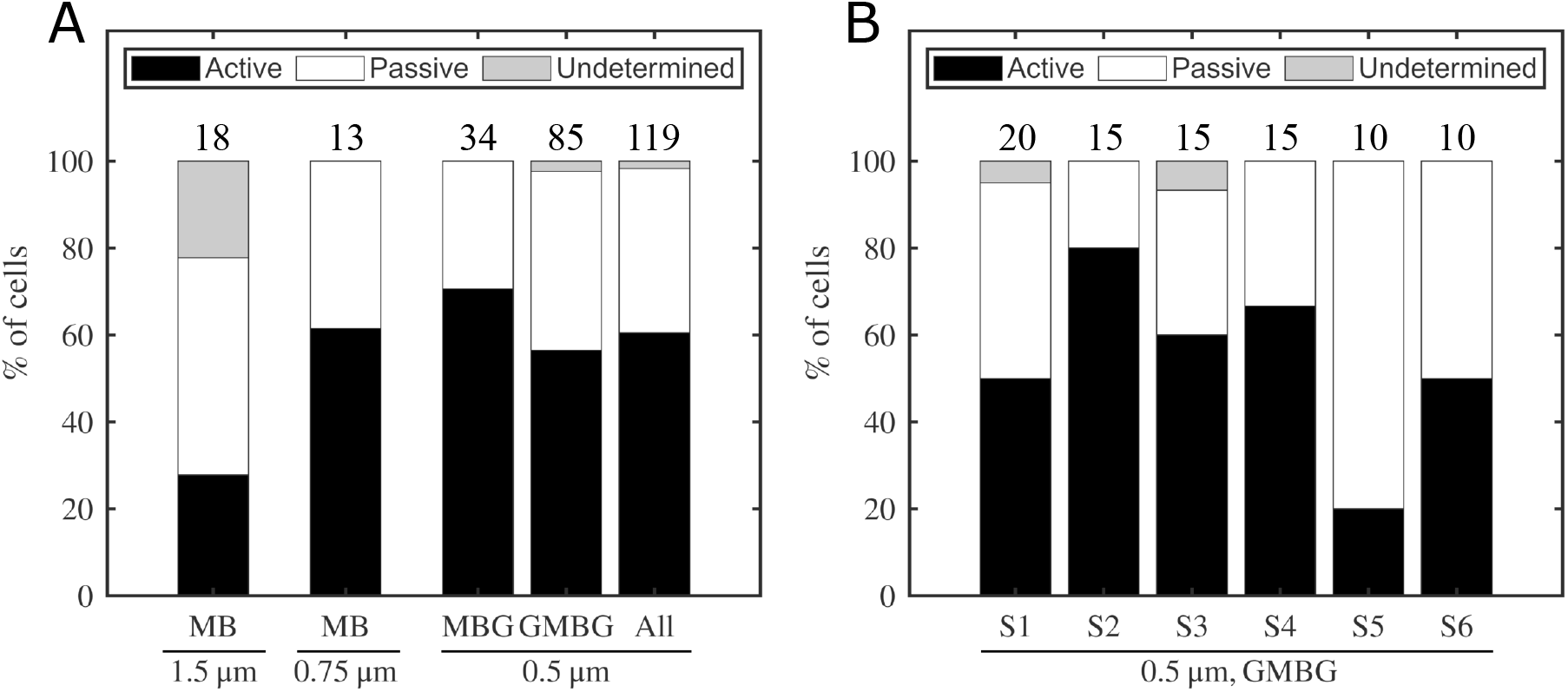
Proportion of active, passive and undetermined cell body rotations observed in the geometry of [29]. **A**. Measurements were conducted using EK01 Δ*cheY* cells with 1.5 µm, 0.75 µm, and 0.5 µm beads, in MB (1.5 µm, 0.75 µm beads) and MBG and GMBG (0.5 µm beads), see *Materials and Methods* and Table SI 7. In all conditions, at least 25% of stably rotating cells were passively counter-rotating. **B**. Proportion of active, passive and undetermined cell rotations for each tunnel slide prepared to collect the 0.5 µm GMBG data in panel A. The fraction of passive counter-rotations varied between slides, from 20% to 80%. Numbers displayed above the bars indicate the number of cells measured in each condition. The numbers observed are in line with those reported in Gabel’s thesis [58], where 13 tethered cells reported in the subsequent publication [29] showed linear PMF-speed relationship, and at least 5, which are reported only in the thesis, were deemed “poorly behaved”. At least two of these cells demonstrate signature of the PMF-speed proportionality saturation we observe in this work.

**Figure SI 16.**
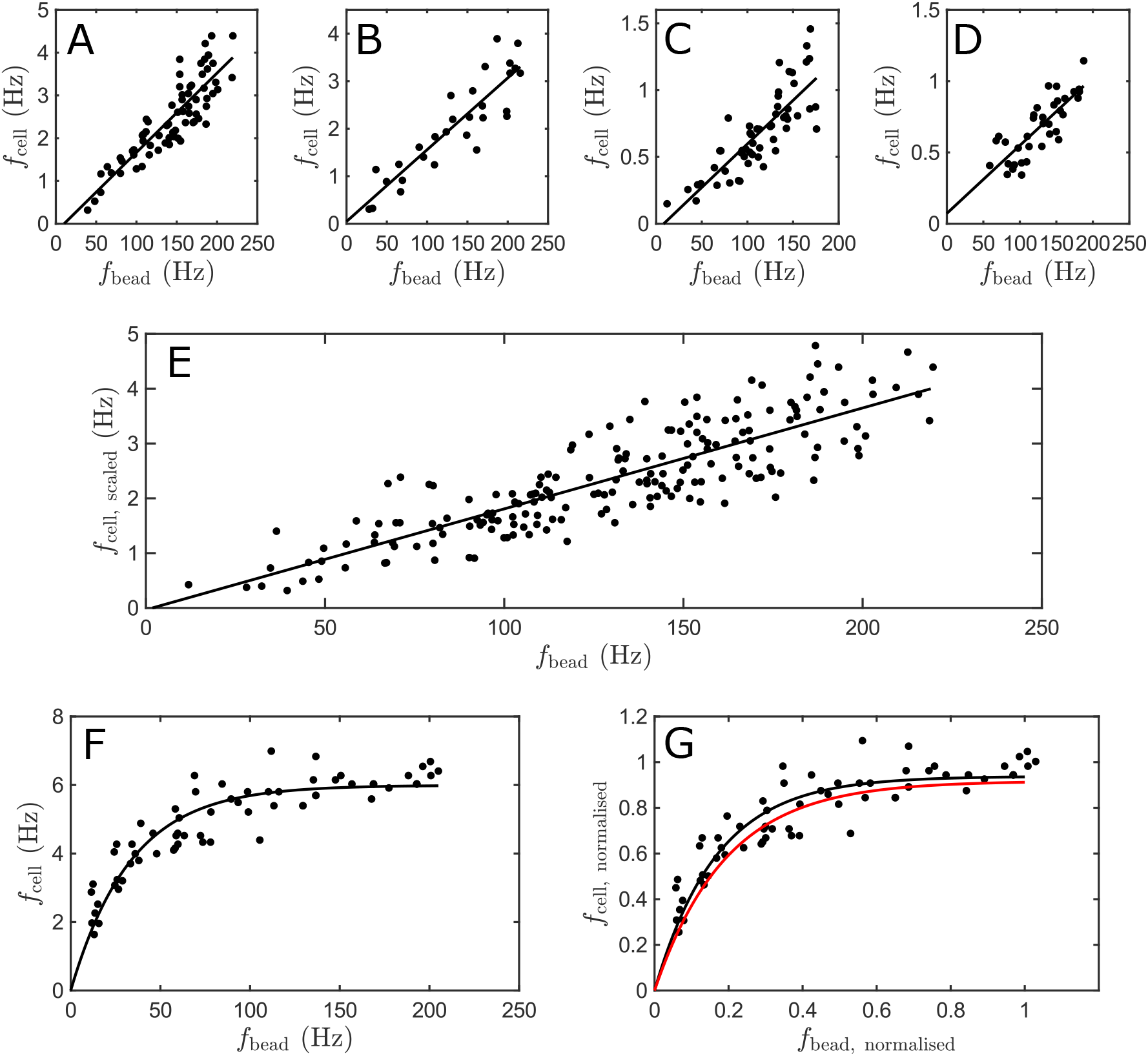
Cell body rotation frequency as a function of bead rotation frequency, for cells exposed to CCCP in the geometry of [29] (see *Materials and Methods*). Passively counter-rotating cells follow the linear relationship reported in [29], whereas an actively rotating cell shows the non-linear relationship of Fig. 2. **A-D**. Traces of 4 passively counter-rotating cells. Unconstrained affine fits yield small intercepts, consistent with the expected linear dependency. **E**. Data of panels A-D replotted on a single graph after rescaling cell body frequencies as in [29]. For each cell, the body frequency is divided by the slope of its fit in A-D, and multiplied by the slope of the fit in panel A, effectively renormalising the body friction coefficient. This produces a single linear relationship between body and bead frequencies, with an affine fit intersecting the vertical axis near the origin. **F**. An actively rotating cell exposed to CCCP shows a non-linear relationship between body and bead frequencies. The black line is a fit using the same function as in Fig. 2. **G**. Data from panel F, replotted after normalising body and bead frequencies (*Materials and Methods*). The black line is the best fit obtained from the equation of Fig. 2. The red line is the fitted master curve of Fig. 2, showing excellent agreement between our bead assay and measurements conducted in the geometry of [29] when both the cell and bead rotate actively. Cells of panels A, B and F were measured in MB+0.3% glucose. Cells of panels C and D were measured in GMB+0.3% glucose.

**Figure SI 17.**
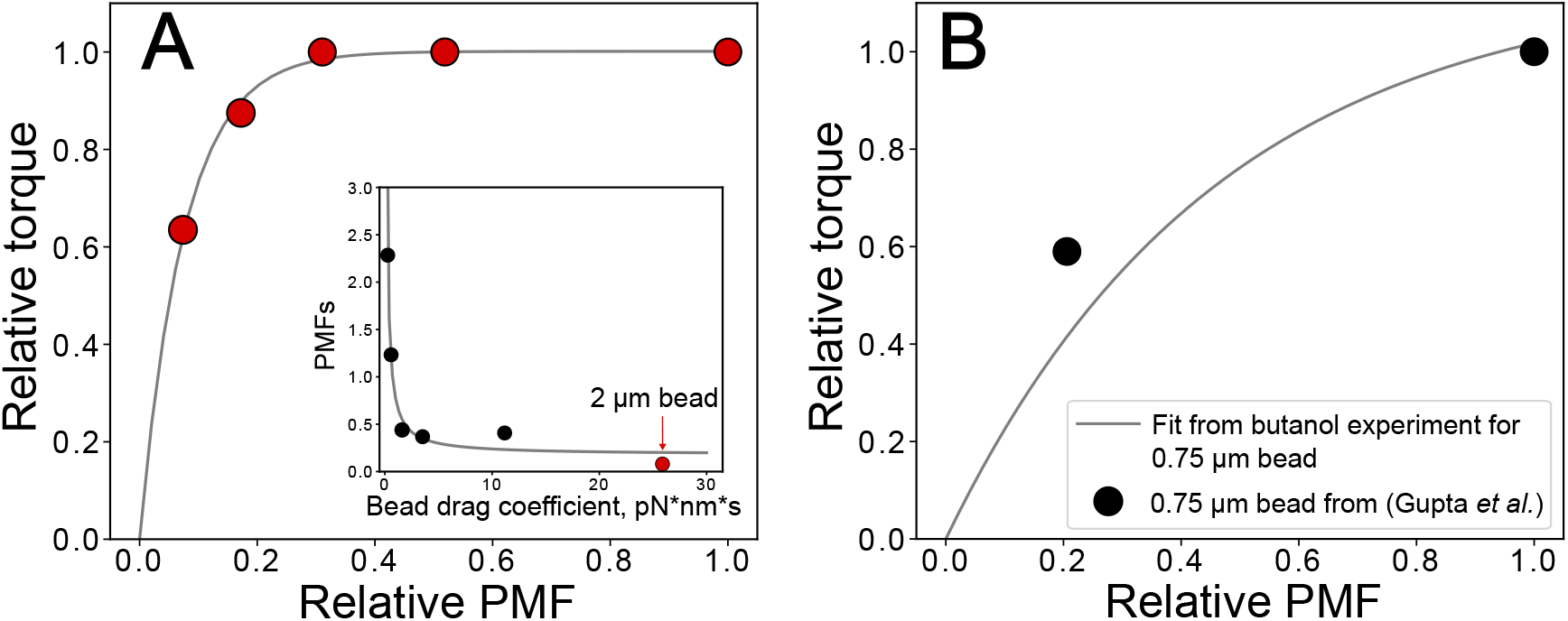
PMF-torque saturation in Gupta *et al*. A. PMF-torque saturation for 2 µm bead from previous work is consistent with this work. The speed data are taken from Gupta *et al*. for cells treated with 0, 0.5, 1, 2, and 5 mM indole [65]. Indole concentrations are converted to relative PMF similarly to butanol, based on the calibration curve in [39], see *Materials and Methods*. The maximum torque, calculated for a 2 µm bead rotating at 10 Hz with 200 nm rotation radius in aqueous environment, is 1630 pN nm, and so within the range we estimated for maximum torque (1108-1839 pN nm). The curve shows the exponential fit as in Fig. 3B: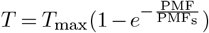, where *T*_max_=1630 pN nm, and PMF_s_ = 0.08. Torque is normalized to the value at PMF=1, which corresponds to cells in MM9+0.3% glucose with no indole added. (***Inset***) PMF_s_ for 2 µm bead is added to the plot from Fig. 3C, and the hyperbolic curve (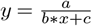, where *a* = 781.7, *b* = 473.5, and *c* = 15.47 are free parameters) is fitted to all the data points. **B**. PMF-torque saturation for 0.75 µm bead from Gupta *et al*. is consistent with this work. The speed data for CCW rotation of the 0.75 µm bead for cells treated with 0 or 2 mM indole are taken from [65], and plotted on top of the PMF-torque curve for the 0.75 µm bead from Fig. 3B. See *Materials and Methods* for normalisation procedures.

**Figure SI 18.**
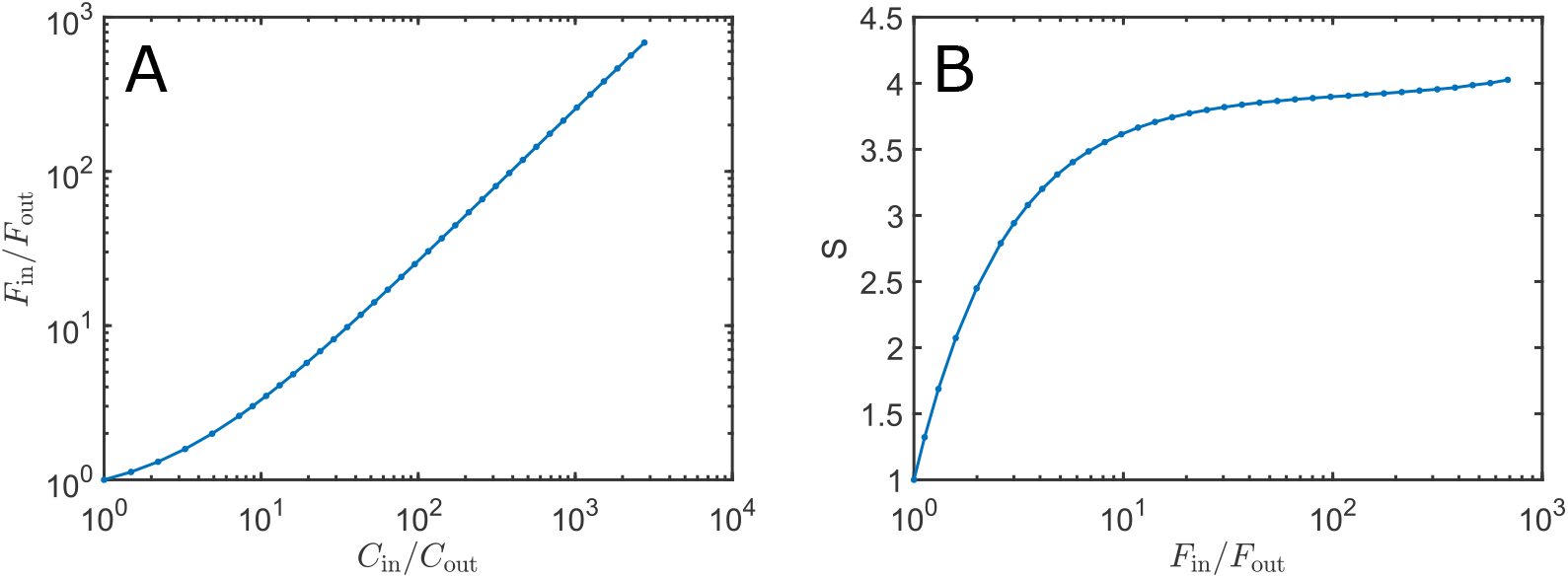
Point spread function correction for TMRM measurements. **A**: Internal/external cell fluorescence ratio measured for known internal/external TMRM concentration ratios in simulated images. **B**: Correction factor *S* as a function of the internal/external fluorescence ratio. See *Materials and Methods* for details.

**Figure SI 19.**
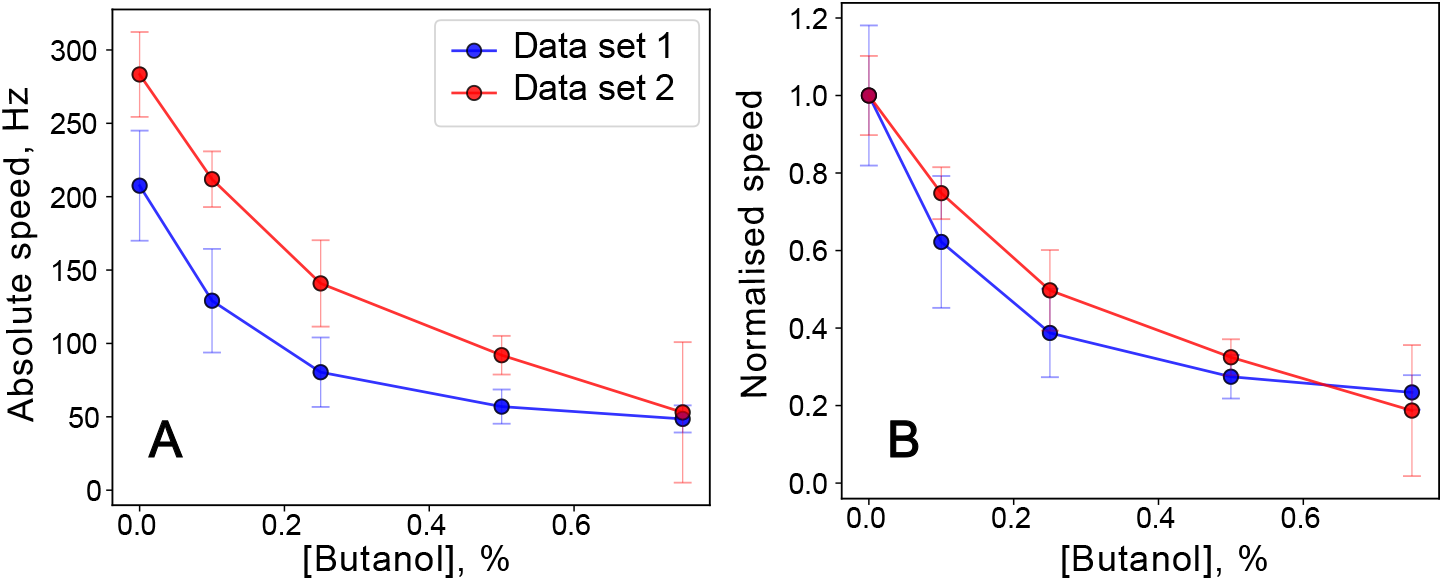
Absolute motor speed can vary between the data sets. (A)An example of two data sets collected with 0.5 µm beads that are particularly different in absolute numbers. Despite the difference, the shape of the butanol-speed curves remains unchanged once speeds are normalised by those measured at 0% butanol in each data set (B). Data points show mean with the tandard deviation.

**Figure SI 20.**
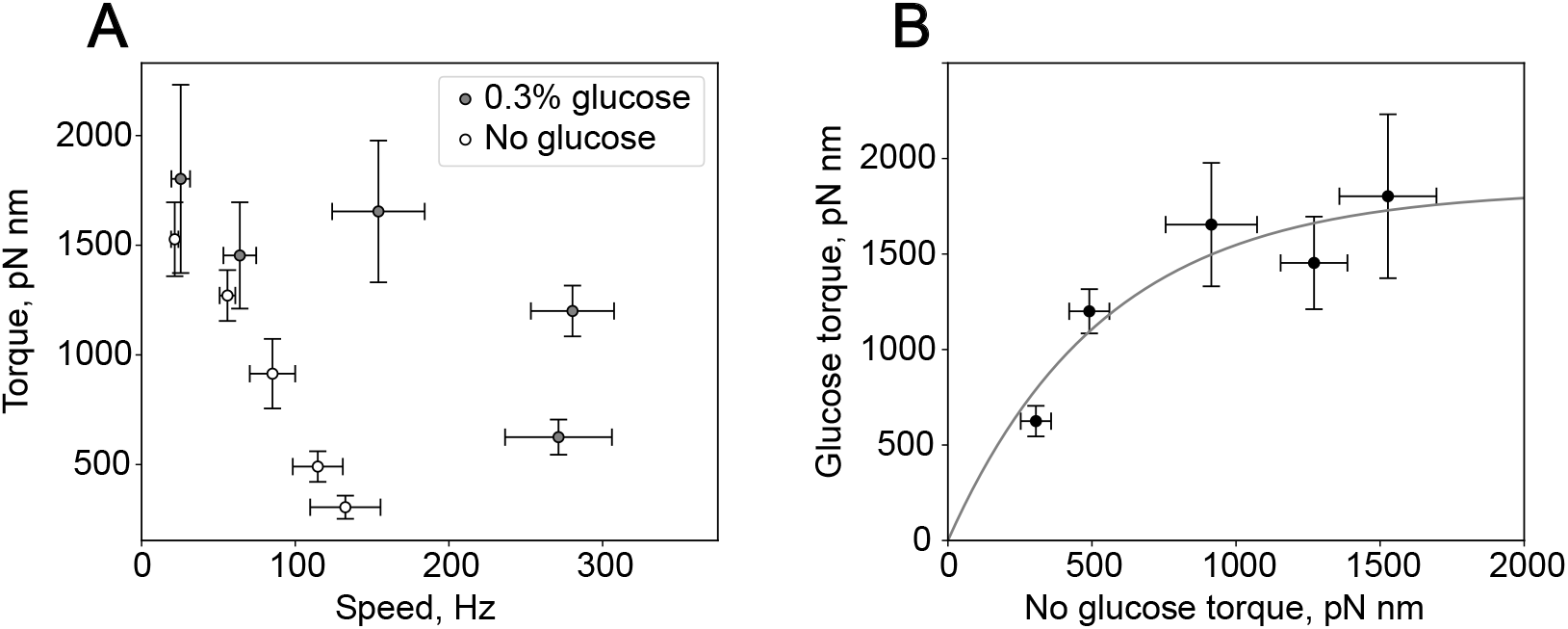
*T*_max_ estimation from the glucose data set in Fig. 1A and SI 1. A. Torque-speed curves for the data set in Fig. 1A and SI 1. **B**. Torque values in the presence of glucose are plotted as a function of torque values in the absence of glucose, for all bead sizes. The line shows the result of an exponential fit, using 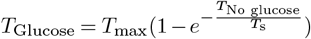 with *T*_max_ and *T*_s_ as free parameters, and returning *T*_max_ = 1839 *±* 247 pN nm. Data points depict mean values with standard deviations, see Table SI 1 for sample size.

**Figure SI 21.**
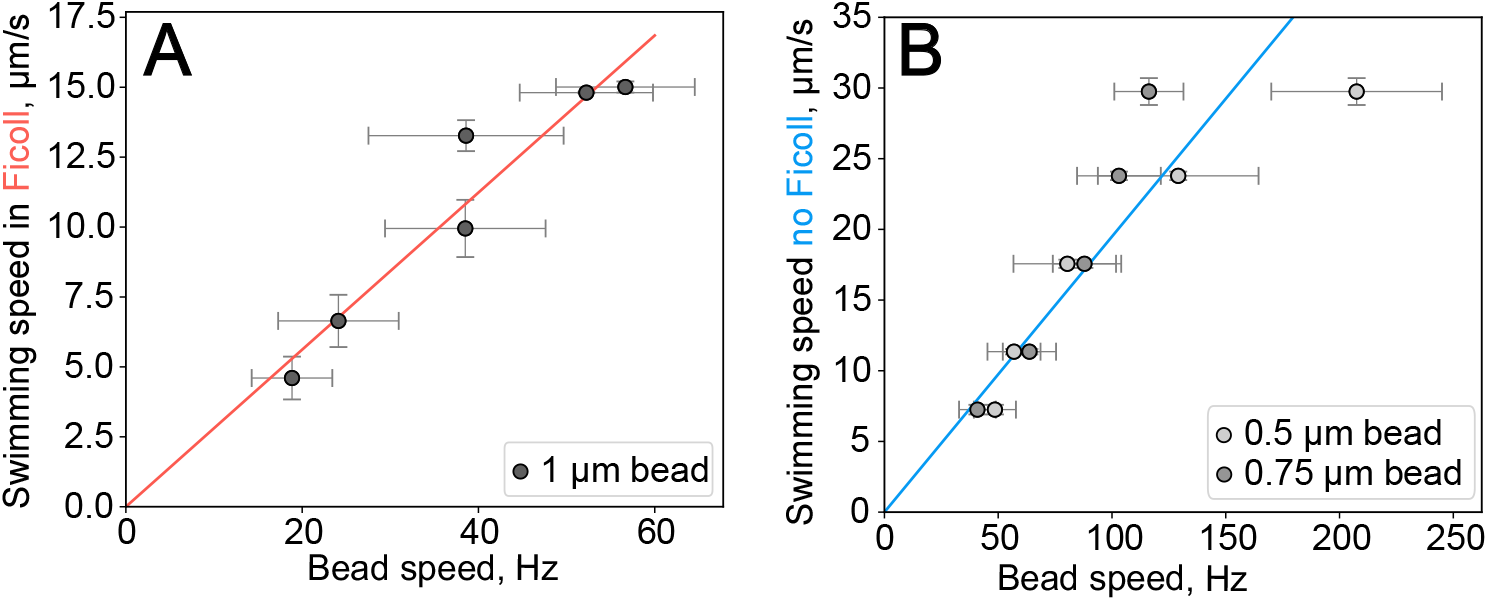
**A**. Swimming speed in MM9+0.3% glucose with 8% Ficoll plotted against the motor speed loaded with 1 µm bead, at the corresponding butanol concentrations. The two speeds are linearly proportional (pink line *y* = 0.28*x*), although he torque is saturated, which indicates that the viscous loads on the motors are equivalent. **B**. Swimming speed in MM9+0.3% glucose media plotted against the motor speed loaded with 0.5 or 0.75 µm bead, at the corresponding butanol concentrations. Neither of the two viscous loads is fully linear with the swimming speed (blue line:linear fit *y* = 0.19*x* applied to all data points) ndicating that the equivalent load lies between the two, consistent with our 0.6 µm bead estimate. Bead colour coding is the ame as in the main text for both (A) and (B). Data points show mean with the standard deviation.

**Figure SI 22.**
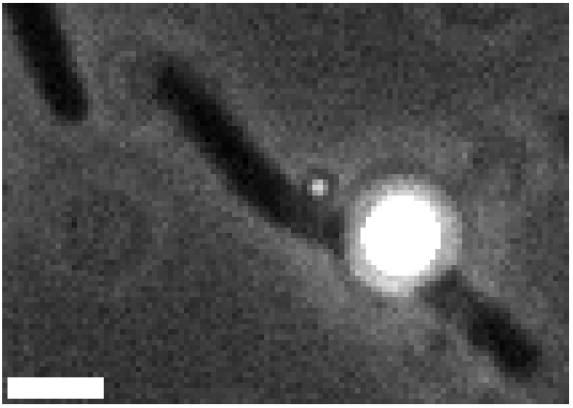
SI Video 1. A cephalexin-treated cell in MM9+0.3% glucose with two beads (0.5 µm and 1.5 µm), each attached to a flagellar motor. The speed of the small bead was measured using this video, and the speed of the large bead was measured using a second video recorded immediately after this one, in which the large bead is in focus. The small bead rotates at 219 Hz *±* 19 Hz, and the large bead rotates at 19 Hz *±* 2 Hz. Video was slowed down 40x. Scale bar: 2 µm.

**Figure SI 23.**
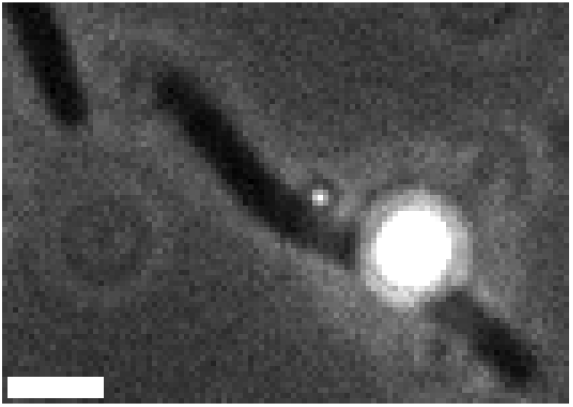
SI Video 2. The same cell as in Video 1, now exposed to 0.5% butanol in MM9+0.3% glucose. The speed of the small bead was measured using this video, and the speed of the large bead was measured using a second video recorded immediately after this one, in which the large bead is in focus. The small bead rotates at 93 Hz *±* 3 Hz, and the large bead rotates at 19 Hz *±* 1 Hz. Video was slowed down 40x. Scale bar: 2 µm.

**Figure SI 24.**
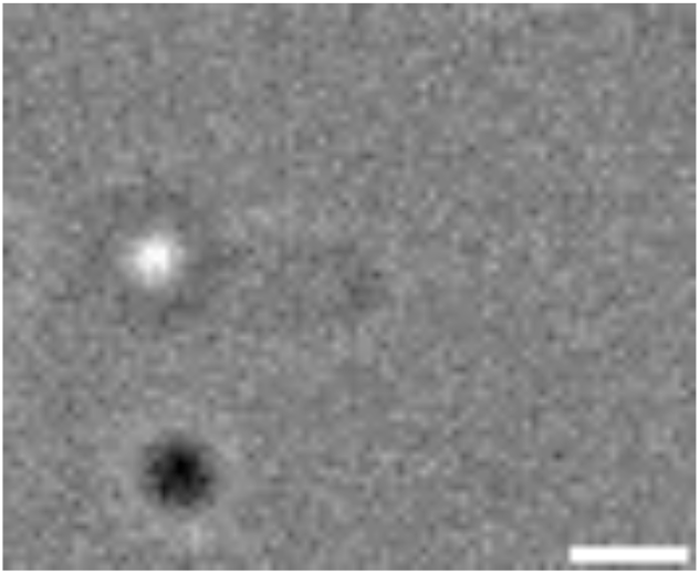
SI Video 3. An EK01 Δ*cheY* cell rotating clockwise, while a 0.5 µm bead attached to it rotates clockwise. In our imaging conditions, this corresponds to both motors rotating counterclockwise when observed from outside the cell. All ecorded tethered cells without a bead rotate clockwise in the field of view. All attached beads rotate clockwise. This indicates hat the rotation of the cell body in this video is an active rotation driven by a functional motor. Video is a 1.6 s-long extract of a 10 s-long video and was slowed down 20x. Scale bar: 1 µm.

**Figure SI 25.**
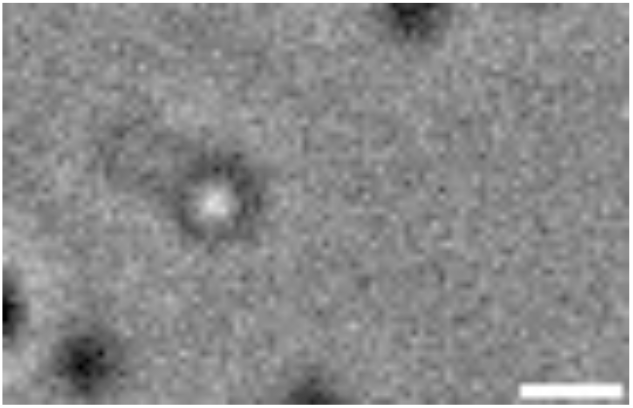
SI Video 4. An EK01 Δ*cheY* cell rotating counterclockwise while a 0.5 µm bead attached to it rotates clockwise. All tethered cells without a bead rotate exclusively clockwise. All attached beads rotate clockwise. This indicates that the rotation of the cell body in the video is a passive counter-rotation induced by the bead rotation. Video is a 1.6 s-long extract of a 10 s-long video and was slowed down 20x. Scale bar: 1 µm.

**Figure SI 26.**
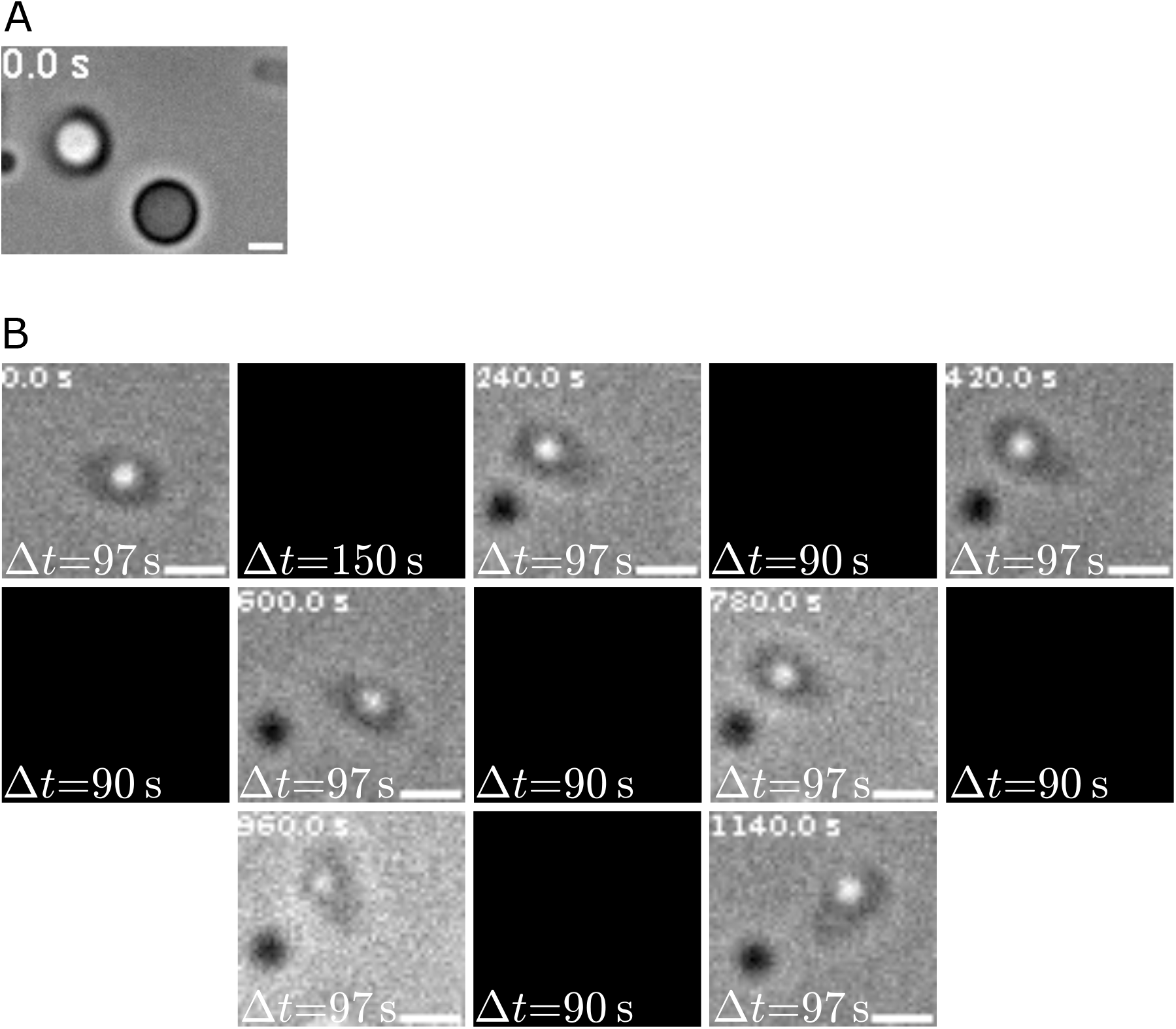
SI Videos 5-8. A. First frame of SI Video 5, a 15 min continuous recording of an EK01 Δ*cheY* cell stably rotating CCW at 3.5-4.5 Hz while a 1.5 µm bead attached to it rotates CW. SI Video 5 is played at true speed. SI Video 6 is identical to SI Video 5, except that it is slowed 10 *×* to visualise the rotation of the bead. **B**. Frames of SI Video 7, a 20 min discontinuous recording of an EK01 Δ*cheY* cell stably rotating CCW at 3-4 Hz while a 0.5 µm bead attached to it rotates CW. Black squares correspond to pauses in the recording between consecutive videos. SI Video 7 was produced by assembling 7 shorter videos (120,000 frames, 97 s each) recorded every 3 to 4 min. SI Video 7 is played at true speed. SI Video 8 is identical to SI Video 7, except that it is slowed down 20 *×*. Whereas both cell bodies rotate CCW in SI Videos 5-8, all tethered cells without a bead rotate exclusively CW, and all attached beads rotate CW, as expected for a Δ*cheY* strain. Thus, stable rotations of the cell bodies in SI Videos 5-8 are passive counter-rotations induced by the beads. Scale bars: 1 µm.

### Supplementary Information Text

#### *V*_m_ *measurements with PROPS*

PROPS measurements confirm our expectations that adding glucose increases the magnitude of the PMF, while adding CCCP or butanol decreases it. To test whether the motor speed is proportional to PMF at low torque, we compared the changes in *V*_m_ and PMF, to the motor speeds measured before and after addition of glucose, Fig. SI 10. For the purpose, we used the PROPS calibration conducted in the original study [43], obtained using induced transmembrane voltage (ITV) and exposure to CCCP. First, we fitted the data of [43] to compute intensity values at all *V*_m_ and intracellular pH pH_i_), Fig. SI 10 A&B. Second, we consider a range of *V*_m_ values around those we measured with TMRM in motility buffer when no glucose is present: - 85 mV ≤ *V*_m,No glucose_ ≤ - 25 mV. Next, we converted them into absolute PROPS intensities using the fitted curve of Fig. SI 10A. Third, we average the final two PROPS intensity measurements of Fig. SI 8 D to obtain 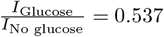. We multiply *V*_m,No Glucose_ in absolute PROPS intensity by this ratio to obtain *V*_m,Glucose_ in absolute PROPS intensity and convert it to *V*_m,Glucose_ using the same calibration curve of Fig. SI 10A. Here, we make a reasonable assumption that the intensities in the previous calibration and in our conditions are linearly related.

Having obtained *V*_m,No glucose_ and *V*_m,Glucose_ we now calculate the corresponding PMF values using:

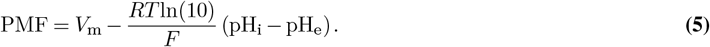

where *R* is the ideal gas constant, *T* the temperature, *F* the Faraday constant, and pH_e_ the extracellular pH. Based on previous measurements, including our own, we take internal pH as pH_i_ = 7.6, both with and without a carbon source [39, 87, 88].Finally, we compute the PMF ratio 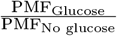and compare it to the low-load motor speed ratios. The calculated PMF atios (1.6 - 2.4) are within error of the recorded speed ratios (1.9 - 2.4), regardless of the value chosen for V_m,No glucose_, demonstrating that the relative motor speed change observed at low load after glucose supplementation can be directly attributed to a similar relative PMF change. We estimated the uncertainty on 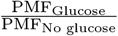 from the standard deviation of the final two PROPS intensity measurements of Fig. SI 8 D, *s* = 0.14. Considering 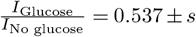 yields, for each *V*_m,No glucose_, a range of *I*_Glucose_ and, thus, of *V*_m,Glucose_. This *V*_m,Glucose_ range is converted into a range of PMF_Glucose_ and finally 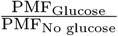, represented by the error bars in Fig. SI 10 C. Additional sources of uncertainty such as the goodness of fit in Fig. SI 10 A, the absence of calibration data below −160 mV, or possible pH_i_ variations, were omitted.

We repeated the above described approach for the control experiment using CCCP, Fig. SI 7. We, thus, estimated that the collapse of *V*_m_ should increase PROPS intensity by a factor 3.1, and collapse of pH_i_ from 7.6 to 7.0, by a factor 1.15. Assuming hat these two effects act independently, the PROPS intensity should increase by a factor of 3.5 upon exposure to CCCP, ∼ 30% higher than the experimentally measured ratio, Fig. SI 7. Here we assumed that the initial *V*_m,Glucose_ in MM9 + glucose is 145 mV [36, 89], and that CCCP sets *V*_m_ = 0 and pH_i_ = pH_e_ = 7.0. The small discrepancy may be linked to inaccuracies ntroduced by the assumptions needed for this analysis, e.g. variations in optical setups and partition of the protein between membrane and cytoplasm can introduce small non-linearities or an increase in pH_i_ upon glucose addition can explain this discrepancy.

Finally, we note that butanol induces smaller intensity changes when compared to CCCP at a concentration for which we expect ull PMF collapse. We observed PROPS aggregation at 1% and 2% butanol in many cells, while GFP remained homogenously distributed, Fig. SI 9. PROPS aggregation appears to be independent of membrane depolarisation, because it did not occur when cells were depolarised with CCCP only (we also note that CCCP is a carrier type protonophore [90], whereas butanol is ikely a pore forming one [38]). Thus, while it qualitatively confirms that butanol dissipates the PMF, we conclude that PROPS cannot be used quantitatively in these conditions.

#### Motor speed variations and maximum torque estimates

The data included in this work were collected in multiple labs by multiple researchers. When accounting for the variations in bead sizes (see Table SI 8), rotation radii (0.1 - 0.2 µm for smaller beads, 0.2-0.3 µm for larger beads), temperature (19-23°C), or culture OD (2.0 - 2.6), there is still some unaccounted variability n measured speeds. We note that when accounting for variability based on temperature differences, we used estimates based on a chimeric motor, where the stator units are powered by sodium [64]. While these differences do not affect our two main conclusions as illustrated in Fig. SI 19, they affect our estimates of the maximum torque, the range of which we find to be 1108-1839 pN nm. This range is obtained by fitting saturating exponential functions to all the data sets. The lowest estimate of *T*_max_ = 1108 *±* 69 pN nm we get from the fit of the 1.5 µm bead butanol data in Fig. 3A, as described in the main text. The highest *T*_max_ = 1839 *±* 247 pN nm is calculated from the glucose data set in Fig. 1A and SI 1. Here, since the full PMF-orque dependence is not available, we plot the torque values calculated in the presence of glucose as a function of the torque values without glucose for all bead sizes, and fit this curve using 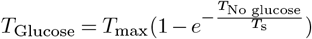 with *T*_max_ and *T*_s_ as free parameters, Fig. SI 20. The thus obtained range of maximum torque values is consistent with previous estimates for the limiting orque of the BFM [12, 26, 34].

#### Viscosity measurements

Diffusion coefficients obtained from DDM analysis for 0.5 µm and 1 µm beads with and without 8% Ficoll (see *Materials and Methods*) were converted to viscosity values using the Stokes–Einstein–Sutherland equation: 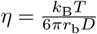, where *η* is the unknown viscosity, *k*_B_ the Boltzmann constant, *T* the temperature, *r*_b_ the bead radius and *D* the diffusion coefficient measured via DDM.

As a control, we first calculated the viscosity of pure water (1 mPas) and obtained *η*_w_ = 0.93 *±* 0.01 mPas and *η*_w_ = 0.85 *± 00*.1 mPas with 0.5 µm and 1 µm beads respectively. In 8% Ficoll, we obtained *η*_F_ = 3.39 *±* 0.02 mPas and *η*_F_ = 2.84 *±* 00.2 mPas with the same bead sizes. Given the systematic deviation from the expected value for water at room temperature, we decided to base our analysis on the relative viscosity increase caused by the addition of 8% Ficoll, the average value of which was 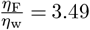when considering both bead sizes.

#### Equivalent load estimates

Since swimming in 8% Ficoll shows a speed response to butanol treatment identical to that of a 1 µm bead in a bead assay conducted without Ficoll, main text Figure 4B and Fig SI 21A, we conclude that the swimming load in 8% Ficoll corresponds to the load produced by the 1 µm in diameter bead (*r*_b,F_ = 0.5 µm). To determine the equivalent bead size of the swimming load in pure medium (no Ficoll) we re-arrange equations 1 and 2 to solve it for *r*_b,w_:

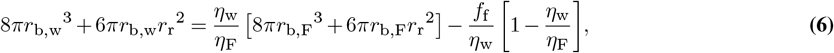

with *f*_f_ the friction coefficient on the filament stub defined in the main text, 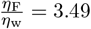 measured in experiments, and *η*_w_ = 0.95 mPas. The result is *d*_b_ = 2*r*_b_ = 0.6 µm, consistent with the experiments main text Figure 4A and Fig. SI 21B.

#### Theoretical speed estimates for passive rotation of tethered cells

For completeness, we theoretically consider the scenario we experimentally confirmed in Fig. SI 16 and SI Videos 3-8, in which the cell is tethered to the slide via a broken motor or a hydrophobic patch on its outer surface [55], thus allowing passive rotation of the cell body. For simplicity, we focus on rotations of the bead and cell body about the same axis, which is experimentally likely since detection of body and bead rotations required alignment with a single pinhole in [29]. Then, the torque-free condition reads *T*_b_ + *T*_cell_ = 0, with *T*_b_ and *T*_cell_ the torques acting on the bead and the cell body, respectively. From this we obtain 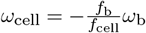, with *ω*_cell_ and *ω*_b_ the respective angular velocities of the cell and bead, and *f*_cell_ and *f*_b_the respective rotational friction coefficients.

To estimate the rotational friction coefficient of cells, we use 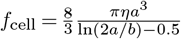 [58], with 2 µm *<* 2*a <* 4 µm and *b* = 0.45 µm [58]. We estimate the friction coefficient of the bead as described in *Materials and Methods* of the main text, with *r*_b_ = 0.2 µm, *r*_r_ = 0.2 µm (lower bound in Gabel and Berg’s experiments [58]), and *f*_f_ = 0.1 pN nm rad^−1^ s.

A passive body counter-rotation speed is then 1-5.5 Hz for a bead rotating at 100 Hz and 2-11 Hz for a bead rotating at 200 Hz, depending on cell size, and in agreement with the values observed in [29] and Fig. SI 16. And, the speeds will vary linearly with the small bead rotation speed as per the torque balance equation.

In the estimates above we neglected any additional friction at the tethering point, and the proximity of the glass slide, as we do not think these affect our estimates (whose upper bounds are already higher than the measured frequencies).

#### Differences between the command voltage and the voltage experienced by the motor in Fung and Berg [24] experiment

Command voltage applied in the Fung and Berg experiment was between 0 and −200 mV. Accounting for the voltage loss on the pipette, part of the cell inside of the pipette, and the leaky seal between the cell and the pipette tip, the voltage experienced by the motor was calculated to be in the range between 0 to −150 mV. However, it was assumed that applying a voltage across the entire cell is equivalent to the membrane potential, i.e. charge separation in the few nanometer layer close to the inner membrane [42], and thus PMF that the BFM uses. *E. coli* is a gram-negative bacterium, whose inner membrane is shielded from the external environment by the periplasmic space, the cell wall, and the outer membrane [91]. The conductivities of the inner and outer membranes of *E. coli*, estimated with electrorotation, are 0.5-5 and 5-25 µS/cm, respectively [92]. Thus, from 2% to 50% of the potential applied by the microelectrode could drop at the outer membrane, and the maximum voltage on the inner membrane experienced by the motor would be lower (in absolute values) than reported. Depending on which end of the range is taken into account, this would result in either 0 to −75, or 0 to −147 mV range. The 2-50% potential loss estimate includes only the outer membrane and does not consider the conductivities of the cell wall and of the periplasmic space, which may act as a short circuit and lower the voltage applied to the motor even more in [24].

